# *RootSlice* – a novel functional-structural model for root anatomical phenotypes

**DOI:** 10.1101/2022.06.29.498145

**Authors:** Jagdeep Singh Sidhu, Ishan Ajmera, Sankalp Arya, Jonathan P. Lynch

## Abstract

Root anatomy is an important determinant of root metabolic costs, soil exploration, and soil resource capture. Root anatomy varies substantially within and among plant species. *RootSlice* is a multicellular functional-structural model of root anatomy developed to facilitate the analysis and understanding of root anatomical phenotypes. *RootSlice* can capture phenotypically accurate root anatomy in three dimensions of different root classes and developmental zones, of both monocotyledonous and dicotyledonous species. Several case studies are presented illustrating the capabilities of the model. For maize nodal roots, the model illustrated the role of vacuole expansion in cell elongation; and confirmed the individual and synergistic role of increasing root cortical aerenchyma and reducing the number of cortical cell files in reducing root metabolic costs. Integration of *RootSlice* for different root zones as the temporal properties of the nodal roots in the whole-plant and soil model *OpenSimRoot/maize* enabled the multiscale evaluation of root anatomical phenotypes, highlighting the role of aerenchyma formation in enhancing the utility of cortical cell files for improving plant performance over varying soil nitrogen supply. Such integrative *in silico* approaches present avenues for exploring the fitness landscape of root anatomical phenotypes.

**Summary statement:** Root anatomy remains an underutilized target for crop breeding. *RootSlice*, a multicellular functional-structural model of root anatomy, simulates the costs and benefits of diverse root anatomical phenotypes to estimate their utility for plant fitness in unfavorable soil environments.

## INTRODUCTION

Plants employ an underground network of roots to gain anchorage, forage and acquire soil resources, synthesize organic compounds, store photosynthates and nutrients, and interact with the soil microbiome (McNear, 2013). This system of roots exists in various shapes and sizes, comprising different classes and orders of roots. Plant root systems vary with species, genotype, age, pedo-climatic conditions, and availability of soil resources, thus displaying a vast phenotypic diversity from organ to cellular scale (Raven & Edwards, 2001). Each root extends along its longitudinal axis only from its tip region, which consists of four partially overlapping functional zones, namely root cap, meristematic zone, elongation zone, and differentiation or maturation zone (Dolan *et al*., 1993; Kozlova *et al*., 2020). At the root tip, the cap senses gravity, assists soil penetration, and protects the meristematic zone, wherein the cells divide and produce all the cells constituting the root (Kumar & Iyer-Pascuzzi, 2020; Kumpf & Nowack, 2015). In the late meristematic zone, the generated cells expand, defining root diameter and moving cells into the elongation zone where the cells rapidly elongate via vacuolization, consequently pushing the root apex further down into the soil (Kaiser & Scheuring, 2020). In the following differentiation zone, the cells are fully differentiated (Dolan *et al*., 1993). This zone also marks the initiation of lateral and higher-order root formation, establishing new axes of root growth and the development of the root system (Moreno-Ortega *et al*., 2017; Verbelen *et al*., 2006). Mature root tissue may continue to develop through such processes as the formation of cortical aerenchyma (Drew *et al*., 2000), cortical senescence (Schneider & Lynch, 2018), and secondary growth (Strock & Lynch, 2020).

On the radial axis, a matured root is arranged in concentric layers of tissues, which have specific forms and functions. The outermost layer is the epidermis followed by the exodermis, sclerenchyma, cortex, endodermis, pericycle, and stele. The latter consists of parenchyma cells and hosts phloem bundles and xylem vessels as vascular tissues (Dolan *et al*., 1993; Lucas *et al*., 2013). Plasmodesmata connect the cytoplasm of adjacent cells, both longitudinally and radially, across the root tissue (Zambryski & Crawford, 2000). Besides the cortex and stele, all other tissues are often unilayered. The elongated epidermal cells bear the root hairs. The endodermis and at times, the exodermis, acquire a suberized and lignified hydrophobic band (the Casparian strip in the endodermis) (Geldner, 2013), and the pericycle cells exhibit meristematic ability to initiate lateral root formation (Beeckman & De Smet, 2014; Torres-Martínez *et al*., 2019). The cortex accounts for a large portion of root cross-sectional area in the primary tissue of most taxa and its features vary with the class and age of the root; across genotypes; and over different environments (Lynch *et al*., 2021). The size, shape, number, and organization of vascular vessels in the stele vary with genotypes and plant species, particularly among monocots and dicots species (Scarpella & Meijer, 2004). In general, the stele has an alternating organization of the xylem and phloem vessels, which largely remain in contact with pericycle cells (Tomescu, 2021). Often, the cell walls of the parenchyma tissue occupying the stele and surrounding the vascular bundles are thickened and lignified, forming a pith. In contrast to monocots, dicot roots undergo radial expansion and loss of the epidermis, cortex, and endodermis via secondary growth (Strock & Lynch, 2020). Notably, significant phenotypic variation and plasticity in root anatomy are found in crop species (Lynch *et al*., 2021; Schneider & Lynch, 2020).

The root anatomical phenotype is characterized by a combination of different phenes (phene is to phenotype as a gene is to genotype) (Lynch *et al*., 2014), which often varies along individual root axes. These anatomical phenes perform distinct roles in regulating the metabolic costs of root construction and maintenance, uptake and transport of soil and plant resources, adaptations to edaphic stress of both biotic and abiotic origins, and interactions with the soil microbiome (Galindo-Castañeda *et al*., 2022; Lynch *et al*., 2021). Root anatomical phenes that are known to influence these processes include - cortical aerenchyma, cortical senescence, cortical cell file number, cortical cell size (i.e., width and length), cell wall thickness, xylem size and number, root hair length and density, secondary growth, multiseriate cortical sclerenchyma, and the width of Casparian strip (Lynch *et al*., 2021). A synchronized and complex interplay among these phenes determines the utility of root anatomical phenotypes for plant fitness under optimal and edaphic stress conditions. Physiological phenes such as the expression of resource uptake and flux transporters further regulate the efficiency of anatomical phenes associated with resource dynamics (York *et al*., 2013, 2016). The metabolic cost of soil exploration is an important determinant of the extent of soil exploration and therefore the capture of soil resources (Lynch, 2015). Interaction among multiple root anatomical phenes thus significantly impacts both the structural and functional properties of the root system.

Plant growth is often limited by an array of edaphic factors (Lynch, 2019, 2022). To cope with limiting conditions, several root phenes – from morphological to anatomical to physiological levels, are triggered and/or altered. But such adaptive modifications of root phenes come with significant direct and indirect costs to the plant, including metabolic costs, ecological costs, and trade-offs for resource acquisition (Lynch & Ho, 2005; Lynch, 2007). For example, root cortical aerenchyma improves crop performance under low soil fertility and drought by reducing root metabolic costs, but improved soil exploration comes with the tradeoffs of reduced radial transport, mycorrhizal colonization, and increased risk for biotic stresses (Lynch, 2015). A comprehensive understanding of the costs and benefits underlying specific root anatomical phenotypes thus becomes essential for the informed deployment of root phenes for improving crop performance.

Exploring the impact of root anatomical phenotypes on soil resource capture is important for informing the development of resource-efficient crop varieties. However, the utility of a phene state depends on the environment as well as interactions with other phene states in integrated phenotypes (Ajmera *et al*., 2022; Klein *et al*., 2020; Rangarajan *et al*., 2018). The number of potential combinations involving multiple phene states over multiple environments exceeds the capabilities of empirical research (Rangarajan *et al*., 2022). For example, if each of five anatomical phenes of interest exists in one of five states (e.g., varying proportions of root cortical aerenchyma formation), there exist 5^5^ = 3125 phenotypic combinations that need to be assessed over different environmental conditions. In such cases, *in silico* approaches become extremely useful. Functional-structural modeling is one such approach, wherein the tissue or organ of interest could either be represented as a continuum or multicellular structure, depending on the question of interest. Continuum models like *OpenSimRoot* have been useful in evaluating the utility of several root anatomical phenotypes for soil resource capture and plant carbon economy in several crop species (Lynch *et al*., 2021; Postma *et al*., 2017). But such models lack the precise cellular and multicellular geometries required to evaluate the interaction between anatomical phenes at cellular and tissue scales.

Multicellular plant systems are often represented in two dimensions implementing one of two modeling techniques, namely vertex-based (Alt *et al*., 2017; Nagai & Honda, 2001; Weliky & Oster, 1990;) or Cellular Potts (Glazier & Graner, 1993; Graner & Glazier, 1992). The main differences between these alternatives lie in the geometrical representation of individual cells and the way physical and chemical interactions among cells are handled (Band *et al*., 2012a). Early multicellular models used a structured rectangular grid to depict root topologies (Grieneisen *et al*., 2007; Mähönen *et al*., 2014; Mironova *et al*., 2012), however, recent models use realistic root anatomies, derived either from digitized microscopic images (Band *et al*., 2012a; Mellor *et al*., 2016, 2022; Moore *et al*., 2015, 2017) or idealized using computational tools (Di Mambro *et al*., 2017). Different frameworks to develop multicellular models exist, namely *CellModeller* (Dupuy *et al*., 2008, 2010), *OpenAlea* (Collis *et al*., 2022; Pradal *et al*., 2008), *VirtualPlantTissue* (De Vos *et al*., 2017; Merks *et al*., 2011) and *MECHA* (Couvreur *et al*., 2018). These frameworks use a standard programming language (e.g., java, python, or C++) to couple model components, representing different processes. Overall, several multicellular root anatomical models can be found in the literature, either independent or implementing these frameworks (Band *et al*., 2012a; Hodgman & Ajmera, 2015; Rutten & Ten Tusscher, 2019). Most of these are primarily focused on capturing developmental aspects of root biology including hormone dynamics (Band *et al*., 2012b, 2012c, 2014; Mellor *et al*., 2016, 2020; Xuan *et al*., 2016), vascular patterning (De Rybel *et al*., 2014; el-Showk *et al*., 2015; Muraro *et al*., 2014;) and biomechanics involving lateral root emergence (Fujiwara *et al*., 2021; Péret *et al*., 2013), root growth and bending (Dietrich *et al*., 2017; Fozard *et al*., 2013, 2016; Jensen & Fozard, 2015). To our knowledge, only a few multicellular root anatomy models exist that capture the flux dynamics of water (Couvreur *et al*., 2018; Heymans *et al*., 2021) and phosphorus (Ajmera, 2016) implementing realistic root geometry. Surprisingly, to date, no multicellular functional-structural model exists that captures root metabolic cost and soil resource acquisition by explicitly implementing cellular to supracellular morphometry of the root, neither in two nor in three dimensions.

Here we present a novel functional-structural modeling platform for root anatomy: *RootSlice*, which explicitly employs the multicellular anatomy of a root segment in three dimensions to evaluate the influence of various anatomical phenotypes on its underlying rhizoeconomics. *RootSlice* precisely captured various anatomical phenotypes of monocot and dicot roots, confirmed the utility of root cortical aerenchyma and cortical cell file number in reducing root metabolic cost, and substantiated the importance of vacuolar expansion in cell and in turn root elongation. Notably, *RootSlice* enables the quantification of rhizoeconomic variables, which were implemented in the *OpenSimRoot/maize* model (Postma *et al*., 2017) to evaluate the utility of anatomical phenes in improving root phenotype performance under low nitrogen availability. Such multiscale integrative compatibility opens avenues for investigating relationships among root anatomy, root architecture, and soil resource capture spanning from subcellular to plant assemblage scales.

## MATERIALS AND METHODS

### Model description

#### Background

A novel three-dimensional multicellular model for root anatomy – *RootSlice*, has been developed with the primary objective of quantifying the influence of different anatomical phenes, individually and in combinations, on soil exploration efficiency of the root.

#### Design and Modules

*RootSlice* model is implemented in C++ and utilizes VTK (i.e., visual tool kit, https://vtk.org/) libraries for final image rendering in 3D data viewing platforms such as ParaView (http://www.paraview.org, Ahrens *et al*., 2005). It consists of three modules: 1) Geometry module which defines dimensions and arrangement of ellipsoid/cylindrical cells and subcellular features with x, y, and z radius, 2) Root cortical aerenchyma module which is an extension to the geometry module accounting for the root cortical aerenchyma formation and reduction in metabolic cost of the roots, and 3) Resource module, which estimates nitrogen and phosphorus content (but not their fluxes) and carbon loss in respiration via mitochondria for each cell.

##### i. Geometry module

###### a. Root tissues and volume

The model simulates a stack of multiple radial root cross-section layers, referred to as slices, to capture a three-dimensional root segment of a defined length. Each tissue type in the root segment can have a user-defined variable number of slices. The model develops the tissues of the root segment in a sequential manner starting from the cortex to the endodermis, pericycle, vascular bundles, and stele parenchyma. The vascular bundles in the stele are defined as hollow cylinders. For monocot roots, the defined size and number of metaxylem cylinders are created in a single layer in the stele. The specified number of phloem cylinders of defined dimensions are created in the outermost rings close to the pericycle. The cortex is constructed by adding cortical cell files in a radially outward fashion as described below. Subsequently, a specified number of sclerenchyma cell files and then the outermost layer of the epidermis is overlayed on the developed cortex. The volume of the root segment is computed by integrating the simulated volumes for cell walls, cell membranes, intercellular spaces, and cell organelles corresponding to different tissue types, from the outermost epidermis to the innermost pith (Supplementary Information 1). The model can capture root segments corresponding to different development zones of diverse root classes. The only exception to this is the conical root tip that varies from the cylindrical shape of the other root zones.

The model captures the distinct stele anatomy of monocotyledonous and dicotyledonous roots via a switch in the input file, which directs the implementation of the appropriate stele development. For dicots, the stele is divided into a defined number of abstract concentric layers of specified sizes. A metaxylem cylinder is then created at the center of the stele. Following this, the defined number of metaxylem cylinders of the specified size are arranged in each of these layers moving radially outwards from the center of the stele towards the pericycle. The space surrounding the metaxylem cylinders in each layer is then populated with pith parenchyma cells of defined dimensions. This is done by calculating the inter-radial distance between metaxylem cylinders in each layer. For this, the angle of the arc formed between each set of metaxylem is calculated. If metaxylem cylinders are tangential, then the angle is zero. If there is no metaxylem cylinder in the layer, then the angle is set to 2π radians. In each layer, metaxylem are placed at equidistant arc length. Metaxylem radius is adjusted so that sum of diameters is not greater than the distance to the protoxylem

###### b. Root cortex

The cortex is defined by concentric files of cortical cells (Figure 1). The number of cell files, cells per file, and dimensions (i.e., diameter, and length) of corresponding cells are specified as input parameters to generate the root cortex. Likewise, cell number and size can be specified for the epidermis and endodermis. A parabolic equation is used to describe the variation of cell diameter according to the position of the cell file whereby cellular diameter increases from the innermost to mid-cortical file and then decreases from mid to outermost cortical file (Chimungu *et al*., 2014; Vanhees *et al*., 2020) Supplementary Information 2). As a result, cells of the inner and outermost cortical files are the smallest while the mid-cortical files have larger cells. The multiplier value in the parabolic function can be updated in the model, if needed, for other species. Users can also specify individual cortical cell files by giving a specific cell size for each cell file.

**Figure 1.**
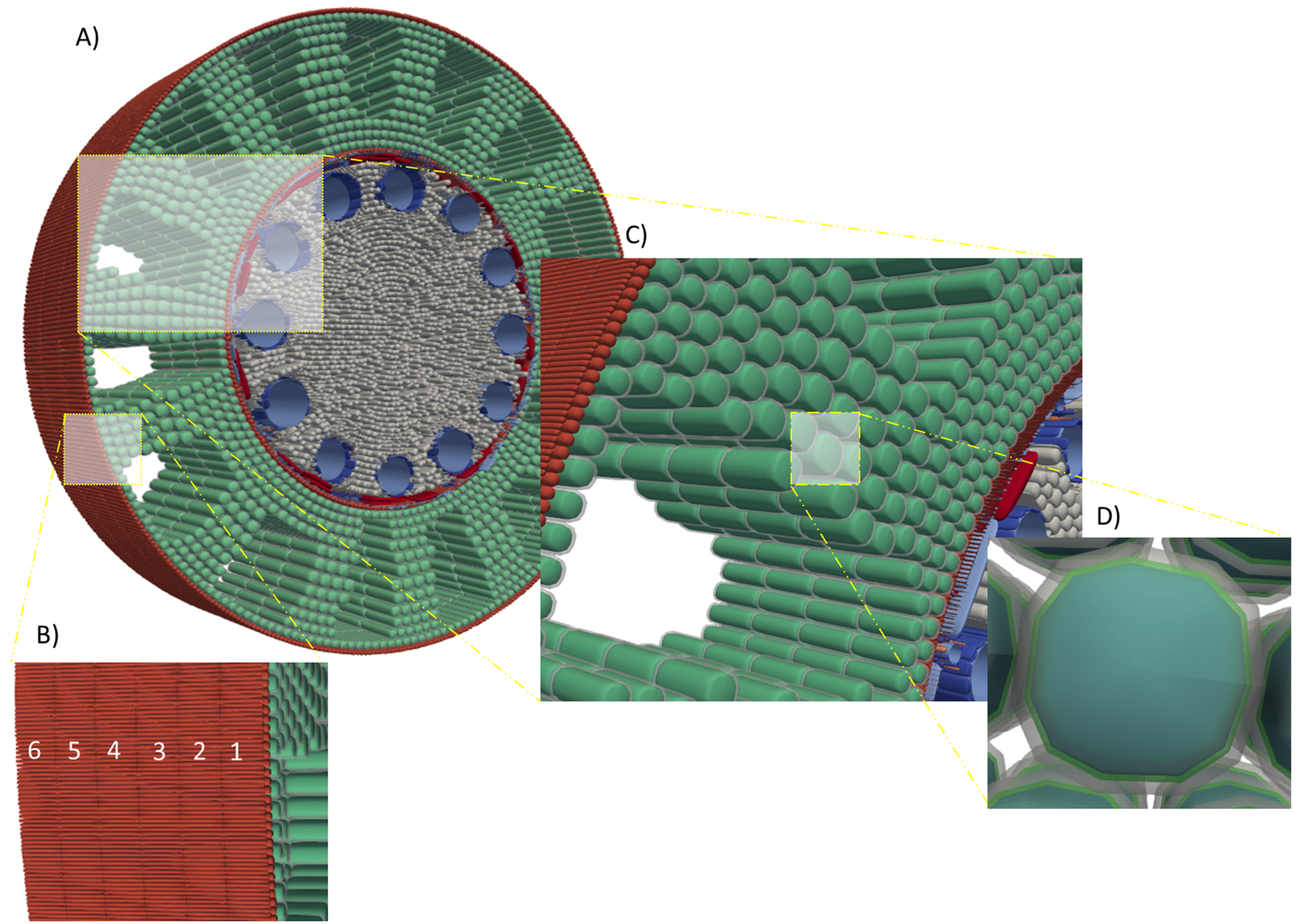
*RootSlice* model showing simulated maize root section. A) Maize root section with eight cortical cell files, stele diameter of 1200 µm, average cortical cell diameter of 60, 12 metaxylem vessels with average metaxylem vessel diameter of 70 µm, and Root Cortical Aerenchyma: cortex ratio of 0.4. B) Close up of longitudinal section showing six cell layers of epidermal cells (numbered 1 to 6). C) Closeup showing cortical cells (green), and aerenchyma (white spaces). D) closeup of a cortical cell showing cell wall (grey), plasma membrane (green), and vacuole (blue).

###### c. Subcellular features

Each cell in the model consists of a cytoplasmic, vacuolar, and apoplastic compartments. The vacuole is represented as a super ellipsoid object located in the center of the cell delineated by the tonoplast. The space surrounding the vacuole forms the cytoplasm (i.e., between the plasma membrane and tonoplast), and the cytoplasmic volume is thus the difference between cellular and vacuolar volume. Currently, the quantified ratio between the cytoplasm and vacuole volume cannot be implemented directly into a three-dimensional model. For this, the distance between the cell membrane and tonoplast is set such that the distance in the X, Y, and Z directions is the same. In this manner, different ratios of cytoplasm and vacuole volume can be simulated.

##### ii. Root cortical Aerenchyma

Programmed cell death of cortical cells leads to the generation of lysigenous root cortical aerenchyma. Different proportions and arrangements of aerenchyma across the cortex can be simulated by defining the input parameters extracted from the root cross-section images. In the model, aerenchyma is represented by removing a certain number and contiguous arrangement of cortical cells leading to the formation of lacunae, while keeping the inner and outermost cortical file intact (Drew *et al*., 2000). Root cortical aerenchyma formation initiates from mid-cortical cells and progresses radially (Drew *et al*., 2000). The potential maximal area of a lacuna is divided into five zones (with reference to the first cortical cell that underwent programmed cell death) and the aerenchyma develops in the following sequence in the module, namely middle, inner, outer, left, and right zones (Supplementary Information 3). In the middle zone, the aerenchyma develops both inwards and outwards cell by cell at first, then from left to right by line (Supplementary Information 3). In the inner and outer zones, the aerenchyma develops cell by cell from the middle to the sides. In the left and right zones, the aerenchyma develops from the middle to both inwards and outwards in the radial direction. The identifier of cortical cells of each lacuna is stored in a database. Cell death in each lacuna stops as soon as the predefined aerenchyma volume is reached. At a certain lacuna number and aerenchyma ratio with a different combination of cell file number and cell size, the gap between lacunae will be different. A search algorithm is used to solve the smallest gap of adjacent lacunae, at the same time, the ratio of the total length of adjacent lacunae’s smallest gap to the perimeter of the middle cortex (rcaGapRatio) is calculated.

##### iii. Resource Module

The resource module is used to calculate the carbon, nitrogen, and phosphorus content of each cortical cell based on the geometry and aerenchyma module.

###### a. Carbon

Root carbon cost refers to the carbon cost of root construction and maintenance. The estimation of root construction involved computing the carbon cost of building the cell wall and cytoplasm comprising root tissue. The average cell wall density was calculated to be 1.51 g cm^-3^, using the cell wall composition of maize roots (i.e., 47 % cellulose, 39 % hemicellulose, 5 % pectin, 5 % lignin, and 2 % protein) (Vatehová *et al*., 2016). Using a cell wall density of 1.51 g cm^-3^ and an average percent carbon mass to be 43.41 %, the mass of carbon needed to build a cell wall was calculated. Based on the glucose equivalent value of cellulose, hemicellulose, lignin, and protein formation (McDermitt & Loomis, 1981), the metabolic cost of building the cell wall was estimated to be 0.48 g cm^-3^. In total, the carbon cost of constructing 1 cm^3^ of the cell wall was estimated to be 1.13 g carbon.

The construction cost of cytoplasm was estimated by using glucose equivalent values (GV) of cytoplasmic macromolecules including carbohydrates, proteins, lipids, amino acids, and nucleic acids (McDermitt & Loomis, 1981). The cytoplasm was estimated to contain 0.075 g cm^-3^ of carbohydrate (Farrar, 1985), 0.081 g cm^-3^ of protein (Brown & Cartwright, 1953), 0.009 g cm^-3^ of lipid (Brouquisse *et al*., 1991), 0.005 g cm^-3^ of amino acids (Brown & Cartwright, 1953) and 0.005 g cm^-3^ of nucleic acids (Brown & Cartwright, 1953; Close & Beadle, 2004). Using the glucose equivalent values for each of these macromolecules, i.e., carbohydrates (0.91), lipids (0.37), proteins (0.48), amino acids (0.46), and nucleic acids (2.27), the carbon cost of constructing cytoplasm was estimated to be 0.025 g carbon per cm^3^ of cytoplasm.

Maintenance respiration values were based on the estimated mitochondrial density per cytoplasmic volume. Mitochondria provide the majority of ATP required by cells (Davies *et al*., 2012; Mathieu *et al*., 1981), so cells with larger numbers of mitochondria will have higher energy demands. The number of mitochondrial per unit volume of cytoplasm can be used to calculate the total number/mass of mitochondria for a given volume of the cytoplasm. The units used for quantifying the mass or number of mitochondria in the literature vary, such as the number of mitochondria per cell, (Avers & King, 1960; Bendich & Gauriloff, 1984) per unit cell area (Griffin *et al*., 2005; Miroslavov & Kravkina, 1991) or per unit volume of cytoplasm, (Robertson *et al*., 1995).

In the model, we use the number of mitochondria per unit volume of cytoplasm to estimate maintenance respiration (Juniper & Clowes, 1965). Also, there is an option for *RootSlice* users to choose the mass or number of mitochondria per cytoplasmic volume. Since respiration is driven by mitochondrial density (Kuang *et al*., 2020), respiration can either be estimated based on mitochondrial density or as respiration per cytoplasmic volume. Respiration per cytoplasmic volume was estimated to be 0.050 to 50 µmol CO_2_ cm^-3^ depending on the root zone. Respiration per cytoplasmic volume was estimated by measuring respiration on the 3^rd^ node of maize root segments (genotype B73) using an infra-red gas analysis (IRGA) system in closed mode (LI-6400XT Portable Photosynthesis System; Strock *et al*., 2018). Total cell number and volume were estimated by measuring longitudinal and cross-sectional root anatomy. The volume of cytoplasm and vacuole was estimated by assuming the distance between the tonoplast and plasma membrane to be a constant of 0.2 µm (Sidhu & Lynch, unpublished data).

###### b. Nitrogen

The concentrations of protein, nucleic acids, amino acids, nitrate, and ammonium in the cytosol and vacuole were used to estimate the total nitrogen content. Total protein concentration in the cytoplasm was estimated to be 0.081 g cm^-3^ (Brown & Cartwright, 1953) and the vacuolar protein pool was estimated to be ranging from 0.001 to 0.011 g cm^-3^ (Belton *et al*., 1985; Mettler & Leonard, 1979; Wagner *et al*., 1981). Assuming an average nitrogen-to-protein content to be 16% w/w (Belton *et al*., 1985), proteinaceous nitrogen was estimated to be 925.2 µmol cm^-3^ in the cytoplasmic pool and 11 to 125.6 µmol cm^-3^ in the vacuolar pool. Nitrogen in nucleic acid, amino acid, and low molecular weight organic compound pools were estimated based on the partitioning of the total nitrogen pool as protein (80%), nucleic acids (5%), amino acids (5%), low molecular weight organic compounds (5%), and soluble nitrogen (5%) (Close & Beadle, 2004; Supplementary Information 1).

Published data regarding the compartmental estimation of nitrate and ammonium concentration is rare and varies depending upon the method used (Miller & Smith, 1996). Here, we used values estimated using double-barreled microelectrodes for cytosolic and vacuolar nitrate concentrations (Miller & Smith, 1996, 2008). For maize root epidermal cells, the estimated mean cytosolic nitrate concentration is 3.1 µmol cm^-3^ and the mean vacuolar concentration is 26 µmol cm^-3^ (Miller & Smith, 1996). Ammonium concentration in cytosol and vacuoles of maize mature roots was estimated using *in-vivo* ^14^N-nuclear magnetic resonance spectroscopy (Lee & Ratcliffe, 1991). Ammonium concentration in the cytosol was estimated to be around 10 µmol m^-3^ and ammonium concentration in the vacuole was estimated to be 15 µmol cm^-3^. The soluble nitrogen content was estimated to be 4 - 10% of the total nitrogen pool (Lee & Ratcliffe, 1991, Supplementary Information 1.).

###### c. Phosphorus

In a mature maize root, cytosolic phosphate concentration is ca. 6.7 µmol cm^-3^ over a range of phosphate availabilities and was reduced to 4.1 µmol cm^-3^ in response to very low phosphate availability (Lee *et al*., 1990). In the same study, the vacuolar Pi concentration ranged from 4.1 - 9.2 µmol cm^-3^ in the roots of phosphate sufficient plants but in response to low phosphate availability, this concentration dropped below 1µmol cm^-3^.

#### Assumptions and parameterization

Model assumptions include a) with changes in different phene states, variables like mitochondrial density per cytoplasm, nitrogen and phosphorus content per cytoplasm and vacuole remain constant b) tonoplast to plasma membrane distance is uniform along the circumference of the vacuole, which is true in the majority of the cases but might be different for the pre-elongation zone (Dünser *et al*., 2019), c) nitrogen and phosphorus content is assumed to be the same in different cell files of the cortex as that is the finest scale of measurement in the literature, however, nutrient concentration may vary depending on the concentric position of the cell files. This can be updated in future versions of *RootSlice* as more precise data become available.

Model parameterization can be conducted for any given species by measuring the anatomical phenotypes from radial and longitudinal sections. Anatomical phenes needed to run the base model include cortical cell file number, cell size by cell files, epidermal cell size, endodermal cell size, stele diameter, number and size of metaxylem and protoxylem vessels, and number and size of phloem elements. Subcellular phenes include the size of the vacuole which can be specified by the tonoplast to plasma membrane distance. Cell wall thickness for each type of cell should be specified and cell wall density can be changed but the default is specified to be 1.51 g cm^-3^. Mitochondrial density per cytoplasm, nitrogen, and phosphorus content per vacuole and cytoplasm should be specified. *RootSlice* can be parameterized using the anatomical data extracted from root cross-section images generated via laser ablation tomography (LAT), manual sections, or other methods. In this work, the *RootSlice* models for maize (*Zea mays* subsp. *mays*), rice (*Oryza sativa* subsp. *indica*), wheat (*Triticum aestivum* subsp. *aestivum*), and common bean (*Phaseolus vulgaris*), were parameterized using the anatomical dataset extracted from LAT of root cross-section images from published studies (Ajmera *et al*., 2022; Lopez-Valdivia *et al*., 2022; Schneider *et al*., 2020, 2021) and pearl millet (*Pennisetum glaucum*) was parameterized based on hand sectioned images (Passot *et al*., 2016).

#### Input file

All the parameters for simulating the root anatomy of interest are defined in the XML format as an input data file. These parameters are often defined as constant variables. However, some parameters namely, base radius, and cortex to aerenchyma ratio are also defined as a range enabling the generation of multiple root anatomies for all the possible combinations of those parameters. The binary files for Windows, Linux, and macOS platforms and the template input file are provided in Supplementary Information 4.

#### Output file

At the start of the *RootSlice* simulation, an output folder is generated using the defined name and data time when the run was initiated. This folder includes several VTP files for visualization, tables in CSV format for all the quantitative information, and text files for raw data to debug or cross-check simulation results. The sample output file corresponding to the template input file (given above) is provided in Supplementary Information 4.

### Simulated scenarios

#### Diverse root anatomies

*RootSlice* model was implemented to simulate diverse root anatomies corresponding to different monocotyledonous and dicotyledonous crop taxa, including rice (*Oryza sativa* subsp. *indica*), wheat (*Triticum aestivum* subsp. *aestivum*), pearl millet (*Pennisetum glaucum*), -, maize (*Zea mays*. subsp. *mays*), and common bean (*Phaseolus vulgaris*); and root classes including nodal, brace, laterals, young and secondary thickened basal root. Parameters used for simulating respective root anatomies can be found in Supplementary Information 5.

#### Cell length and vacuolar expansion in a maize nodal root

Root elongation was depicted by a series of discrete simulations corresponding to the quasi-static snapshots of an elongating root segment. At the start, a root segment from the pre-elongation zone consisting of three slices of root tissues each 75 µm in length was simulated with a stele diameter of 240 µm, an average cortical cell diameter of 10 µm, a cortical cell wall thickness of 0.1 µm, and 8 cortical cell files. In the subsequent set of simulations, the slice and cell length were elevated to 150, 225, and finally to 300 µm, thereby capturing the dynamics of an elongating root. For simplicity, all root tissues were assumed to have uniform cell and slice lengths. Distance between the tonoplast and plasma membrane was set to be constant at 0.2 µm across this set of simulations, highlighting vacuole expansion-driven root elongation.

#### Root cortical aerenchyma in a maize nodal root

For simulating aerenchyma formation we simulated a root with 12 cortical cell files, a stele diameter of 400 µm, an average cortical cell diameter of 37.5 µm, and 10 metaxylem vessels with an average metaxylem vessel diameter of 70 µm (Supplementary Information 1.). Different proportions of root cortical aerenchyma (0 to 40% of the cortex volume) were simulated while keeping other parameters constant. A range of 0 to 40% aerenchyma covers more than the 25^th^ percentile to the 75^th^ percentile of the empirical data (Schneider *et al*., 2020). Note that percent aerenchyma in this work is considered as a percentage of cortex volume.

#### Cortical cell file numbers in a maize nodal root

Roots with varying cortical cell files were simulated. The range of files simulated was from 8 to 20 which represents approximately the 25th percentile and 75th percentile of the empirical data (Schneider *et al*., 2020). In all the simulations, base parameters include a stele diameter of 400 µm, an average cortical cell diameter of 37.5 µm, and 10 metaxylem vessels with an average metaxylem vessel diameter of 70 µm.

#### Combinations of different proportions of aerenchyma formation and the number of cortical cell files in a maize nodal root

Four different phene states of cortical cell file number (i.e., 8,12,16, and 20) were factorially simulated with four different proportions of aerenchyma formation (i.e., 0, 10, 20, and 40 % of the total cortical cross-sectional area). Base parameters for all the simulations include a stele diameter of 400 µm, an average cortical cell diameter of 37.5 µm, and 10 metaxylem vessels with an average metaxylem vessel diameter of 70 µm.

#### Integrating *RootSlice* for different nodal root zones with *OpenSimRoot/Maize*

Different maize nodal root zones i.e., pre-elongation, elongation, late elongation, early mature, and mature zone (Kozlova *et al*., 2020), with four different cortical cell files (i.e., 8, 12, 16, 20) and three levels of maximum aerenchyma formation from tip to the mature root zone (i.e., 0-0, 0-20, 0-40 %, Lenochová *et al*., 2009) were simulated by the *RootSlice* model (Figure 2, Supplementary Information 6). The parameters corresponding to each zone are given in Table 1. Cytoplasmic and vacuolar nitrogen content was set to be 0.0018 and 0.0042 g cm^-3^, respectively. Respiration (i.e., carbon released) per unit of cytoplasmic volume was set at 0.050 µmol cm^-3^ (Supplementary Information 1.). The *RootSlice* output for each of these distinct root zones was incorporated as the temporal parameter for each nodal root in the *OpenSimRoot/maize* model (Postma, *et al*., 2017), to evaluate the influence of varying anatomical phenotypes on plant performance. *OpenSimRoot* is a heuristic modeling platform aimed at testing the adequacy of a hypothesis to understand the observed results. In total, twelve integrated root anatomical phenotypes were evaluated using *OpenSimRoot/maize* over four different soil nitrogen supplies with three replicates (Figure 2). A list of *RootSlice* output parameters is shown in Supplementary Information 6. Interactions between these two phenes in each integrated phenotype for shoot biomass performance under low nitrogen supply were quantified wherein the observed (predicted) shoot biomass response of integrated phenotypes was compared to the expected response derived using the additive null model (Côté *et al*., 2016) and the reference phenotype with 12 cortical cell files and no aerenchyma formation. Individual phene state responses were calculated by subtracting the reference phenotype response from each phene combination response that differed in only one phene state. Summing the corresponding individual phene state responses yielded the expected additive response for various phene combinations. To be comparable with actual responses (i.e., predicted shoot biomass), the obtained expected responses of integrated phenotypes were added to the reference phenotype response. The actual response greater than, equal to, and less than the expected response respectively corresponds to the synergistic, additive, and antagonistic relationship between the number of cortical cell files and the proportion of aerenchyma formation. More details on this can be found (Ajmera et al., 2022; Ma *et al*., 2001).

**Figure 2.**
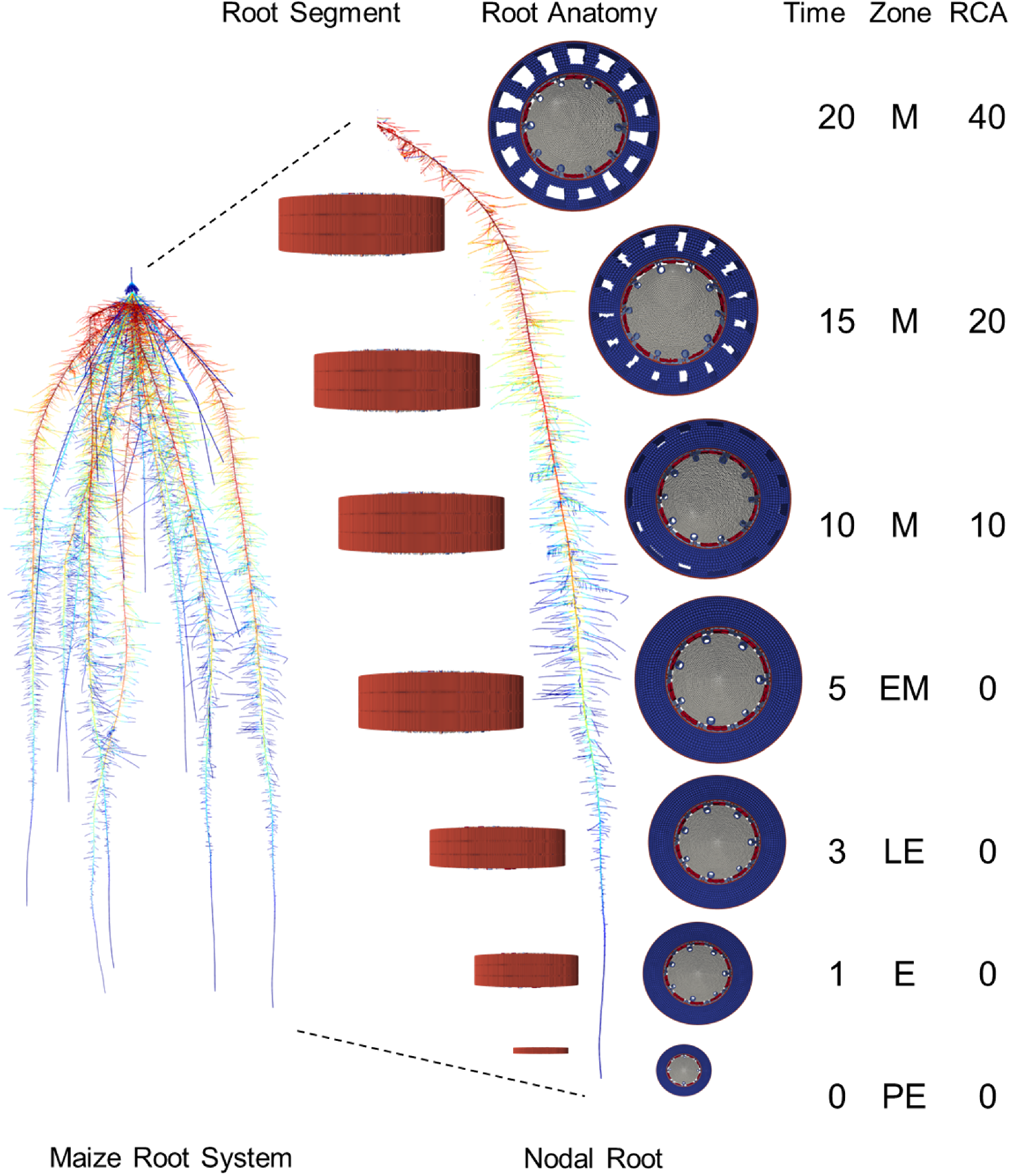
Coupling of *RootSlice/maize* and *OpenSimRoot/maize* models. *RootSlice* models with different root anatomies corresponding to the five root development zones (i.e., PE: pre-elongation, E: elongation, LE: late elongation, EM: early maturation, and M: maturation) of a maize nodal root were simulated. The matured root undergoes three different levels of cortical aerenchyma formation (i.e., 10, 20, and 40 %). In total, seven different root anatomies were simulated in the *RootSlice* model leading to the rhizoeconomic output variables, including root diameter, tissue density, respiration per unit volume, and nitrogen content. Each root zone and corresponding anatomies evolve as the root grows, wherein the cell undergoes the transition from one zone to another. The derived rhizoeconomic variables were temporal scaled with root growth (i.e., transition from one zone to another) and passed to the *OpenSimRoot*/maize model for each nodal root. *OpenSimRoot* simulation of maize root system at 40 days after germination. Color gradient highlights the proportion of root cortical aerenchyma, with red to dark blue respectively denoting 40 to 0% aerenchyma formation.

**Table 1.** *RootSlice* input parameters for different nodal root zones in maize. Elongation zone parameters have been adapted from (Kozlova *et al*., 2020, and Lenochová *et al*., 2009).

| Root zone | Cortical cell wall thickness (μm) | Root cortical Aerenchyma (%) | Cell length (μm) | Cortical cell file number | Stele diameter (μm) | Cortical cell diameter (μm) | Meta xylem number | Meta xylem diameter (μm) |
| --- | --- | --- | --- | --- | --- | --- | --- | --- |
| Mature | 2 | 0, 0.1, 0.2, 0.4 | 100 | 8, 12, 16, 20 | 400 | 30 | 10 | 25 |
| Early mature | 1 | 0 | 100 | 8, 12, 16, 20 | 400 | 30 | 10 | 25 |
| Late elongation | 0.5 | 0 | 75 | 8, 12, 16, 20 | 350 | 26 | 7 | 20 |
| Elongation zone | 0.2 | 0 | 50 | 8, 12, 16, 20 | 300 | 22 | 5 | 15 |
| Pre elongation zone | 0.2 | 0 | 25 | 8, 12, 16, 20 | 250 | 18 | 4 | 12 |

## Results

*RootSlice* correctly simulates longitudinal and radial root anatomy of the distinct root developmental zones – i.e., elongation and maturation, different root classes – i.e., nodal, brace, lateral, and basal and diverse crop species including maize, rice, wheat, pearl millet, and common bean (Figure 3). *RootSlice* also reconstructed the anatomy of an archeological maize root specimen dated to be 5,280 to 4,970 years old (Figure 3D) (Lopez-Valdivia *et al*., 2022). The model captures diverse root anatomies by implementing empirical measurements corresponding to the varying cortical cell file number, cortical cell size, cortical aerenchyma, stele diameter, number, and size of xylem and phloem vessels, and their positions by the surrounding parenchyma cells and pith tissue (Figure 4). This also includes the vascular patterning and secondary thickening distinctive to dicot roots. (Figure 3I, J). For each cell, the model precisely accounts for the volumes of the cell wall, plasma membrane, vacuole, cytoplasm, and protoplasm and thus enabling accurate quantification of the rhizoeconomic variables involving carbon, nitrogen, and phosphorus (Figure 4G).

**Figure 3.**
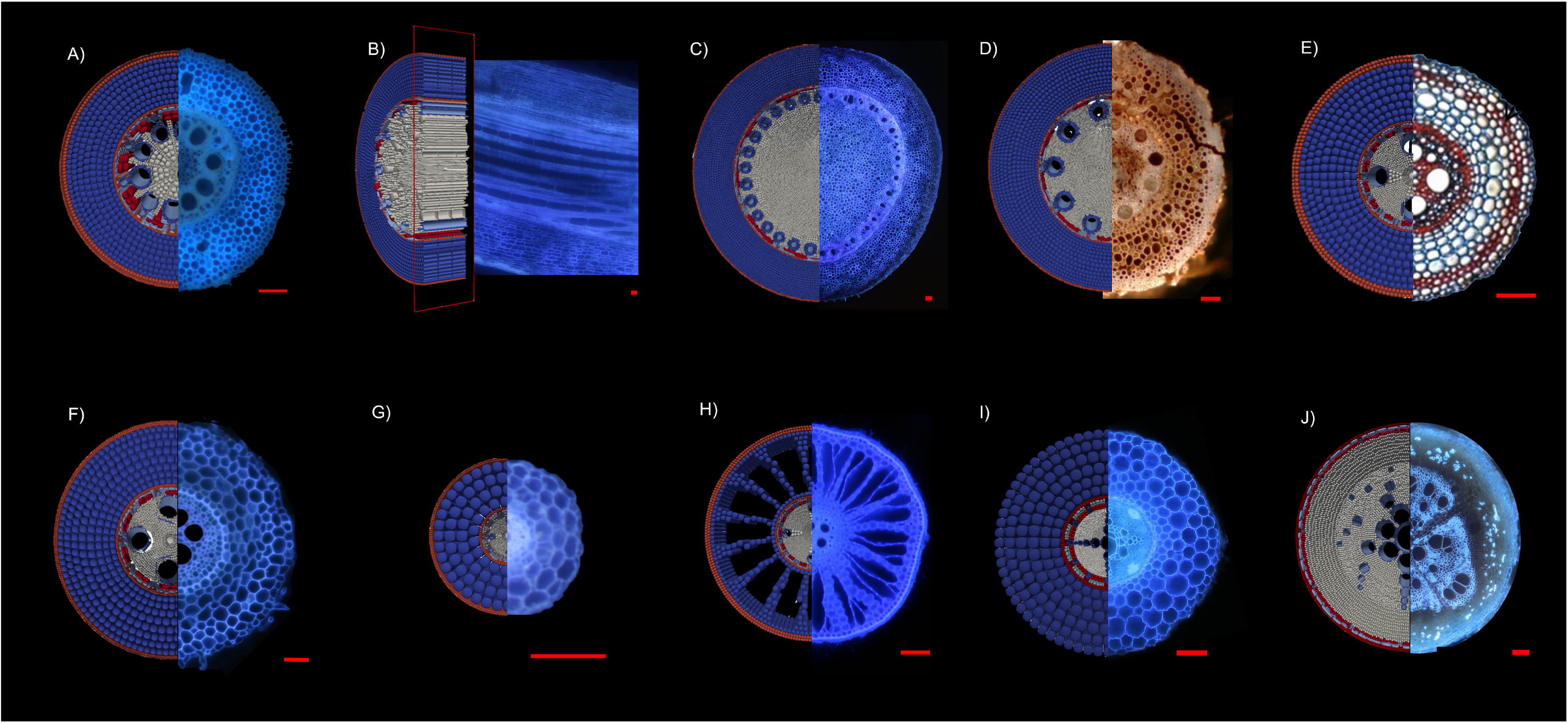
*RootSlice* reconstructions of different root types of a diverse set of crop species. A) Cross-section of maize nodal root, B) longitudinal section of maize nodal root, C) maize brace root, D) ∼5000-year-old maize nodal root, E) pearl millet nodal root reconstructed from a hand section, F) wheat nodal root, G) thin wheat lateral root, H) rice nodal root highlighting aerenchyma, I) common bean basal root and J) secondary growth in common bean. Scale = 100 µm. Note: Digitally ablated longitudinal section of *RootSlice* model compared with laser-ablated longitudinal section.

**Figure 4.**
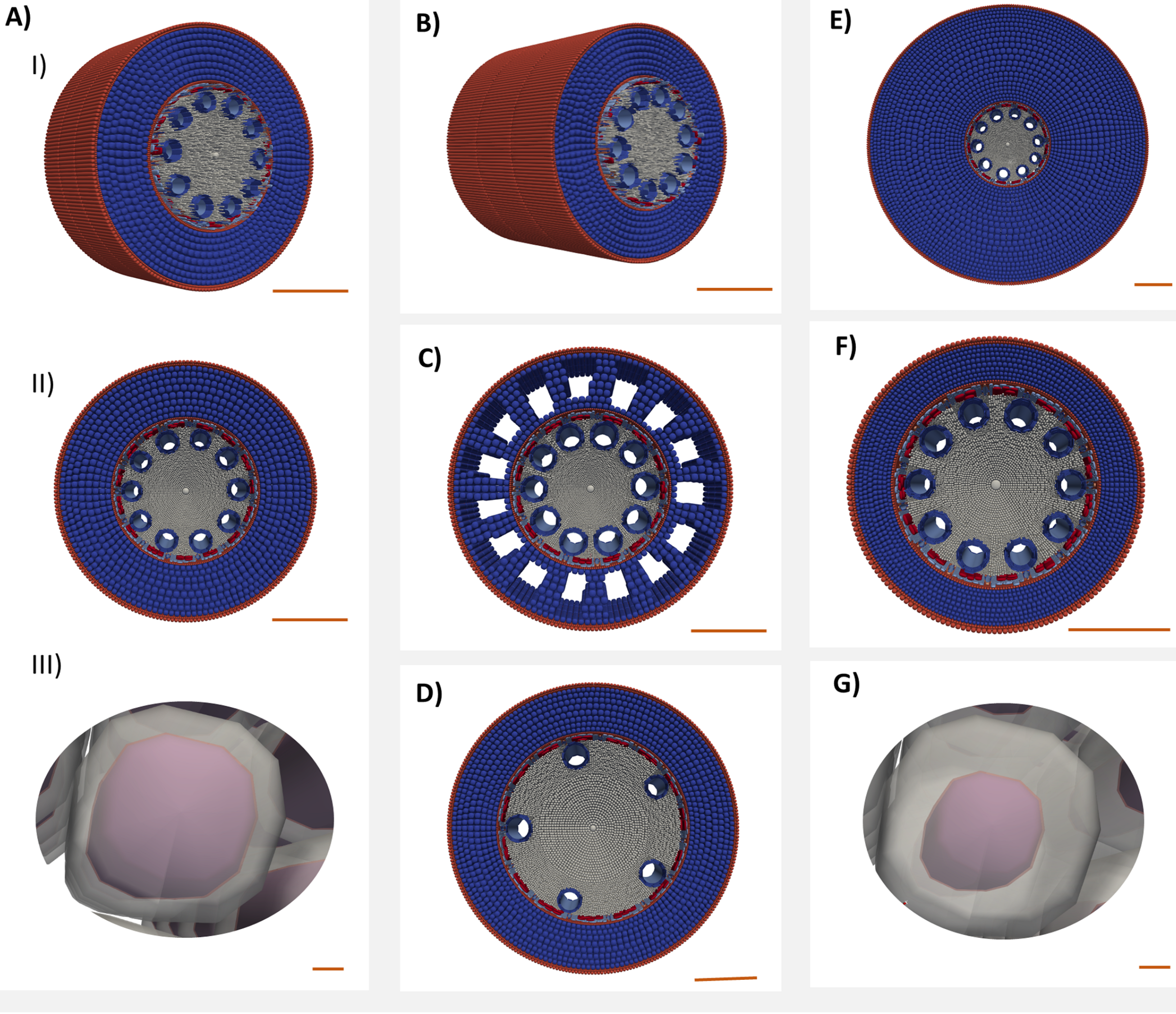
*RootSlice* models showing variation in different maize root anatomical phenes. A. I and A.II) Base model with stele diameter of 400 µm, the average cortical cell size of 30 µm, 8 cortical cell files, 0% root cortical aerenchyma, 10 metaxylem vessels with an average diameter of 25 µm, and cortical cell length of 100 µm, A.III) close of base model cortical cell with 2 µm cell wall thickness. B-G shows single phene variations to the base model. B) Longer cortical cell length of 200 µm, C) 20% root cortical aerenchyma, D) increased stele diameter (600 µm), and few-large metaxylem vessels, E) 20 cortical cell file number, F) smaller cortical cell size (15 µm), G) thicker cortical cell wall (4 µm).

*RootSlice*/*maize* models were created to evaluate three anatomical phenes, namely, vacuole size, root cortical aerenchyma, and cortical cell file number, and the interactions among the latter two phenes on the rhizoeconomy of the plant. These example case studies illustrate the utility of this root anatomy modeling platform.

### Case study 1: Vacuole expansion drives cell elongation

Results show that vacuole occupancy is crucial for cell elongation. Most of the new volume gained in expanding cells is contributed by the vacuole (Figure 5). With every 1 µm increase in cell length, cortical vacuolar volume increases by 94 µm^3^, cytoplasmic volume increases by 7 µm^3^, cell wall volume increases by 3.6 µm^3^, and plasma membrane volume increases by 0.35 µm^3^. Thus, the greatest gain in cell volume (∼90 %) is represented by the expansion of the vacuole, and just < 9 % is attributed to cytoplasmic volume and cell wall volume combined. The carbon cost of constructing cytoplasm is estimated to be 2.51e^-14^ g µm^-3^, therefore with every 1 µm increment in cell length, 1.50e^-9^ g of carbon investment is needed to construct cytoplasmic macromolecules.

**Figure 5.**
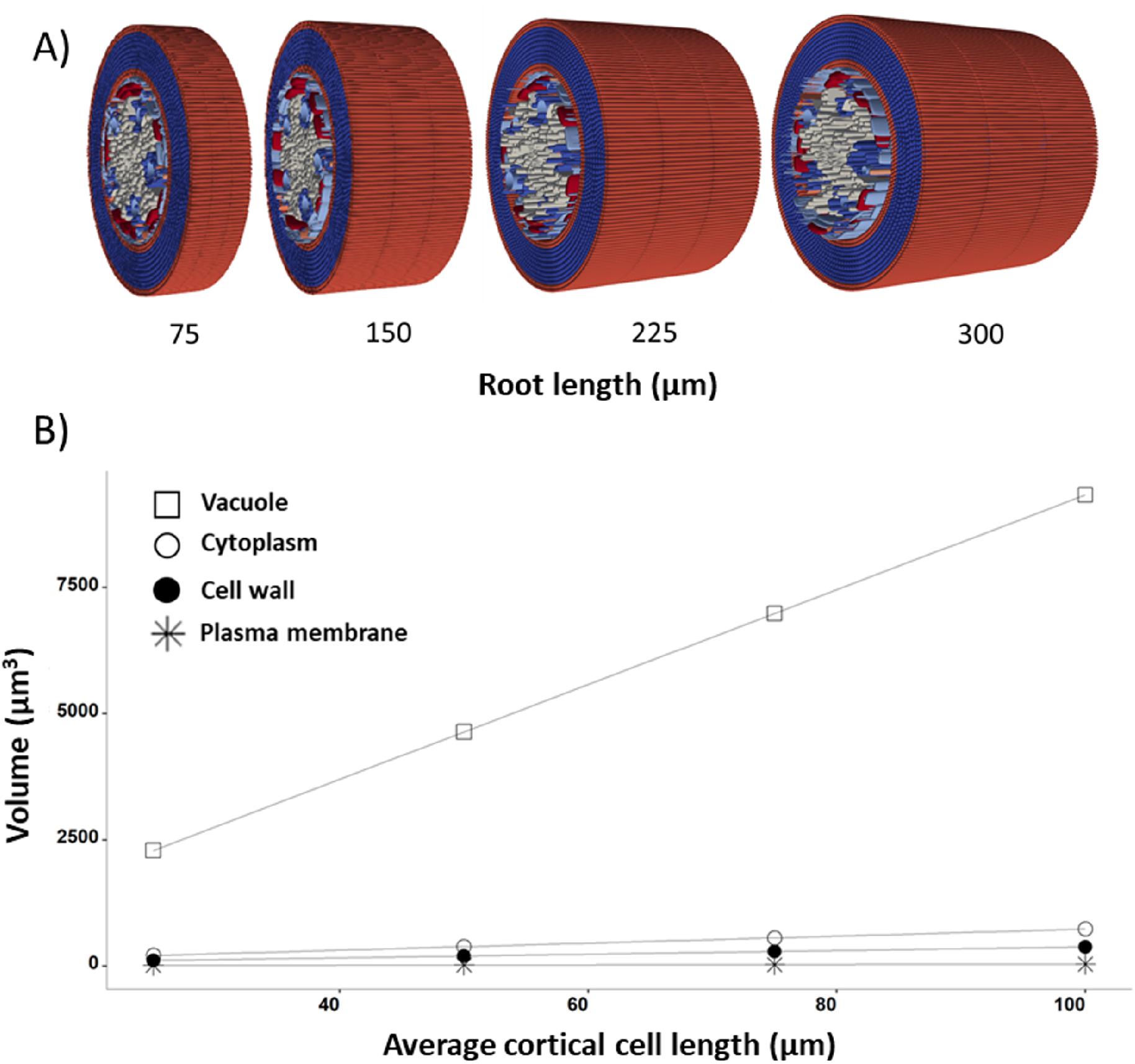
*RootSlice* models showing vacuole-driven cell and root elongation. A) *RootSlice* models show a change in root length going from 75 µm to 300 µm with each root having three cell slices longitudinally, and B) volumetric changes in different cellular compartments and organelles of elongating root cortical cells.

### Case 2: Aerenchyma formation reduces the maintenance cost of the root cortex

For a maize nodal root, increased cortical aerenchyma formation from 0 to 40 % of cortex volume is predicted to cause a 30% decrease in cytoplasmic plus vacuolar volume and a 27% decrease in mitochondrial density per unit cortex volume (Figure 6B, C). Phosphorus content per unit of cortex volume is predicted to decline by ∼30% and nitrogen content by 31 % in response to the increase in aerenchyma formation (Figure 6D, E). Empirical data shows that increased aerenchyma formation from 0 to 20 % reduced nitrogen content ∼25-50 % and reduced respiration rate per unit root length (i.e., including cortex, stele, endodermis, and epidermis) from ∼25-57 % (Figure 7A). In agreement with the empirical data, the model showed a linear decline in respiration rate and nitrogen content per unit of root length with a linear increase in aerenchyma formation (Figure 7B-D).

**Figure 6.**
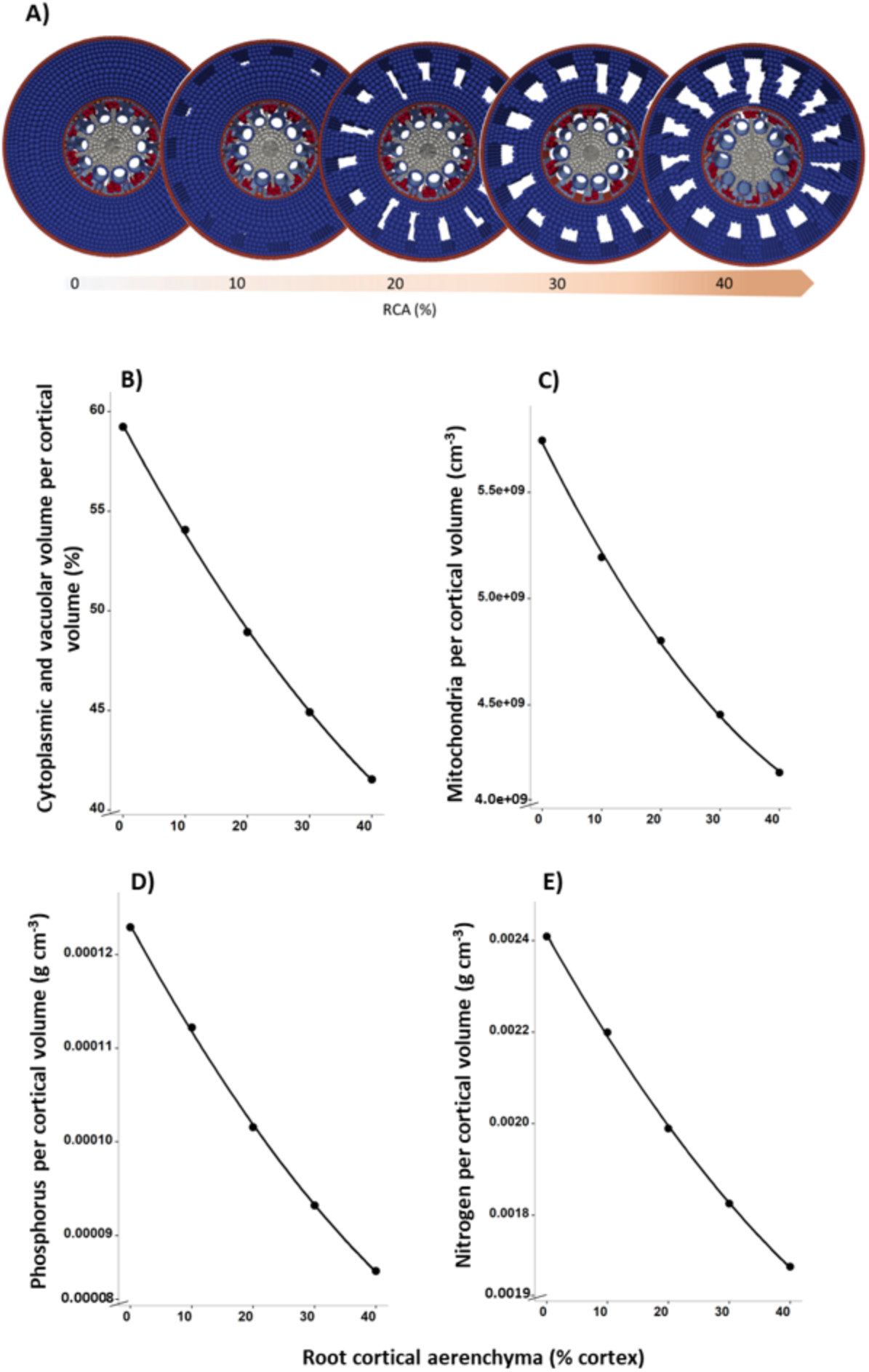
Effect of root cortical aerenchyma on different cellular compartments and resources. A) *RootSlice* models with different proportions of root cortical aerenchyma. B) Effect of root cortical aerenchyma cytoplasmic and vacuolar volume, C) mitochondrial density, D) phosphorus, and E) nitrogen content.

**Figure 7.**
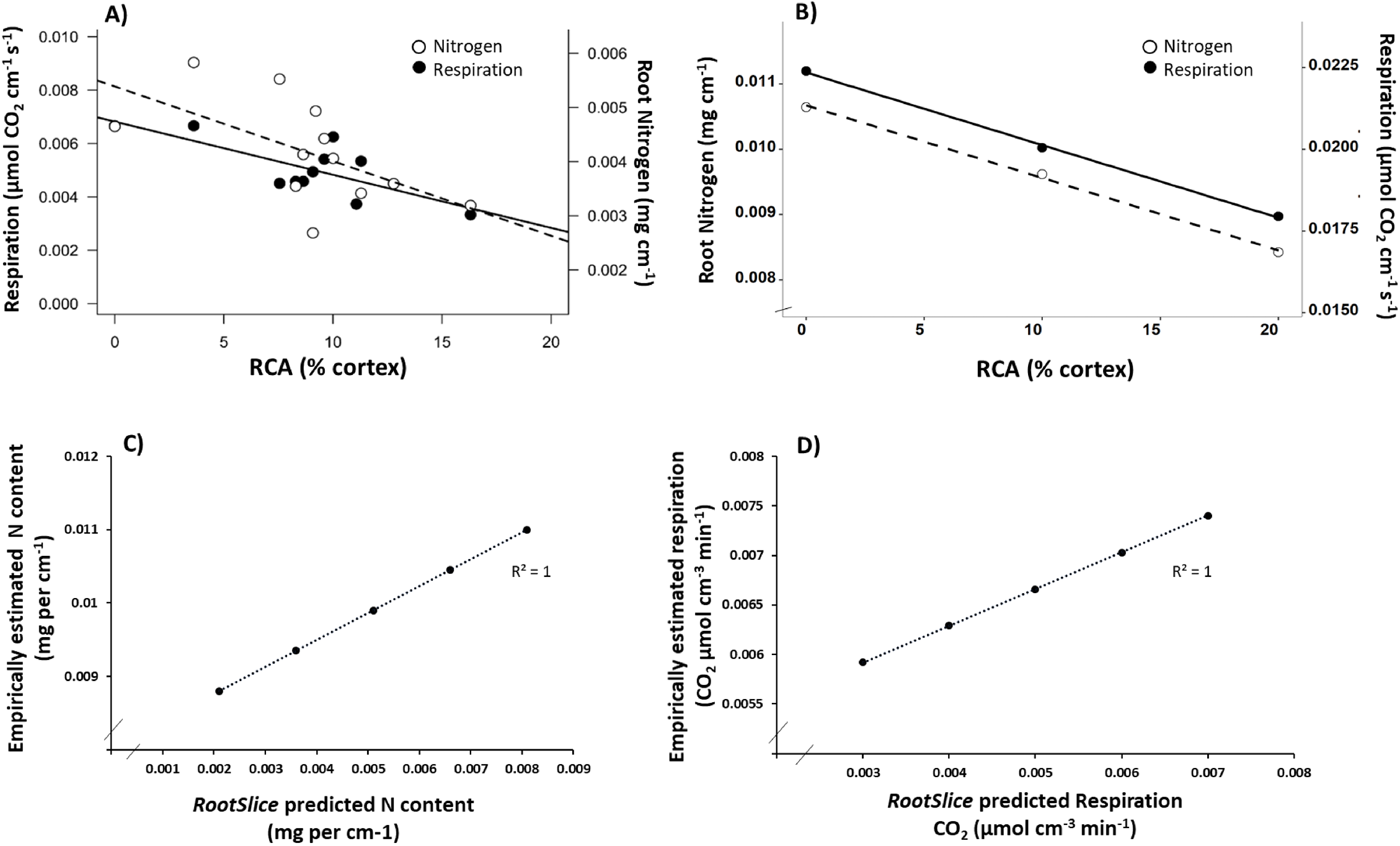
Comparing *RootSlice* predictions about the effect of root cortical aerenchyma on root metabolic cost with empirical data. A) Empirical data showing a decrease in tissue nitrogen content and root respiration with an increase in root cortical aerenchyma (Saengwilai *et al.,* 2014), B) *RootSlice* predicts a decrease in root nitrogen content and mitochondrial density with an increase in root cortical aerenchyma, C) *RootSlice* predicted root nitrogen content vs empirically estimated root nitrogen content, D) *RootSlice* predicted root respiration vs empirically estimated root respiration.

### Case 3: Cortical cell file number increases the metabolic cost of the root cortex

For a maize root, an increase in cortical cell file number from 8 to 20 is predicted to elevate the volume per unit root length of the cell wall by 229% and cytoplasm plus vacuole by 461% (Figure 8B, C). Similarly, the model predicted a 528% increase in the mitochondrial density, a 467 % increase in nitrogen content, and a 453% increase in phosphorus content per unit root length in response to a greater number of cortical cell files (Figure 8C-E). A simulated root segment with 8 cortical cell files is predicted to have 71% less respiration per unit root length than a root segment with 20 cortical cell files. This is in accord with empirically measured respiration of the root segments with 8 cortical cell files being 57-71% less than that with 20 cortical cell files (Figure 9, Chimungu *et al*., 2014). The model predictions thus show good agreement with empirical observations.

**Figure 8.**
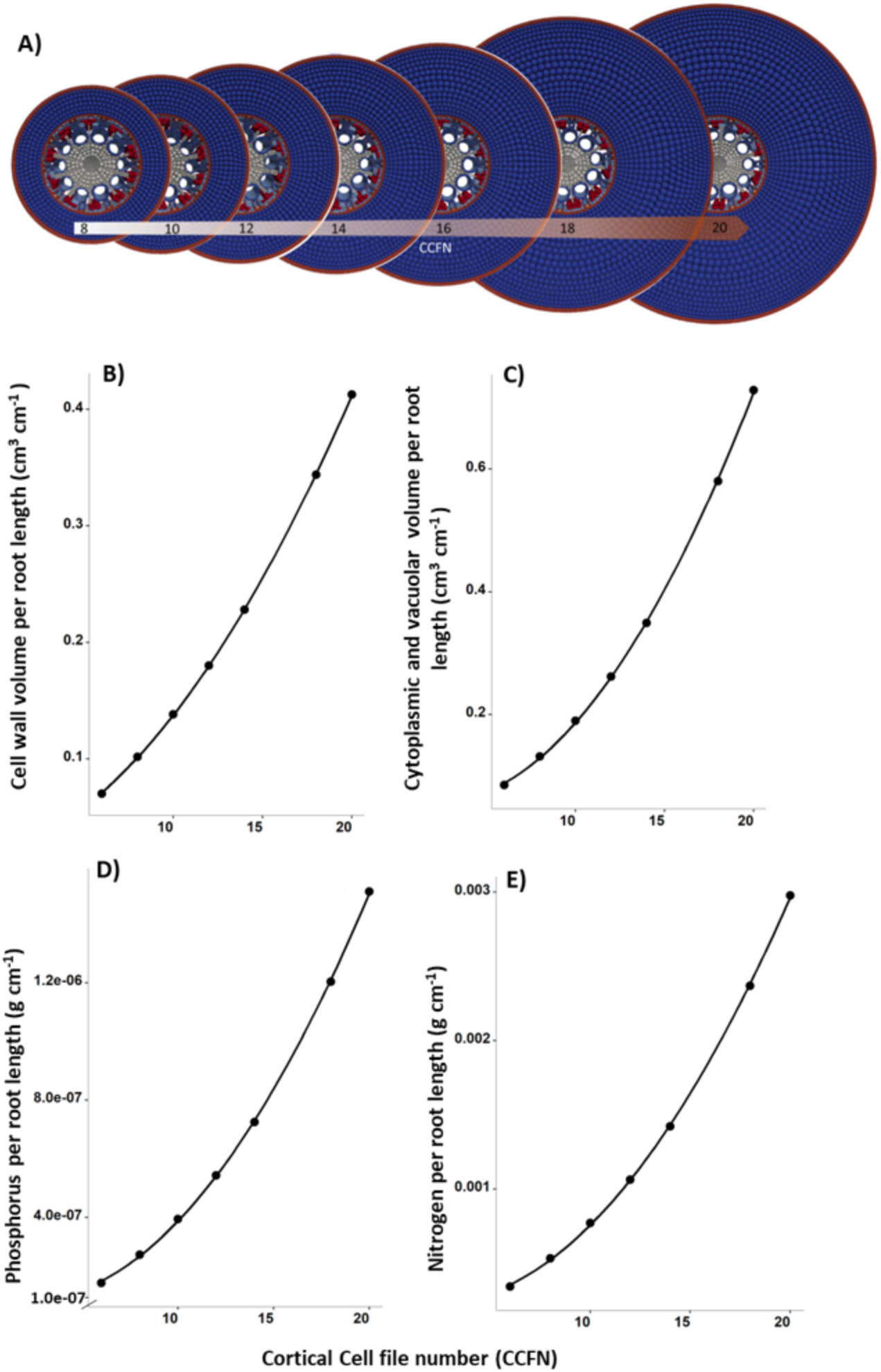
Effect of cortical cell file number on different cellular compartments and resources. A) *RootSlice* models with the varying cortical cell file number. B) Effect of cortical cell file number on cell wall volume, C) cytoplasmic volume and vacuolar volume, D) nitrogen content, and E) phosphorus content.

**Figure 9.**
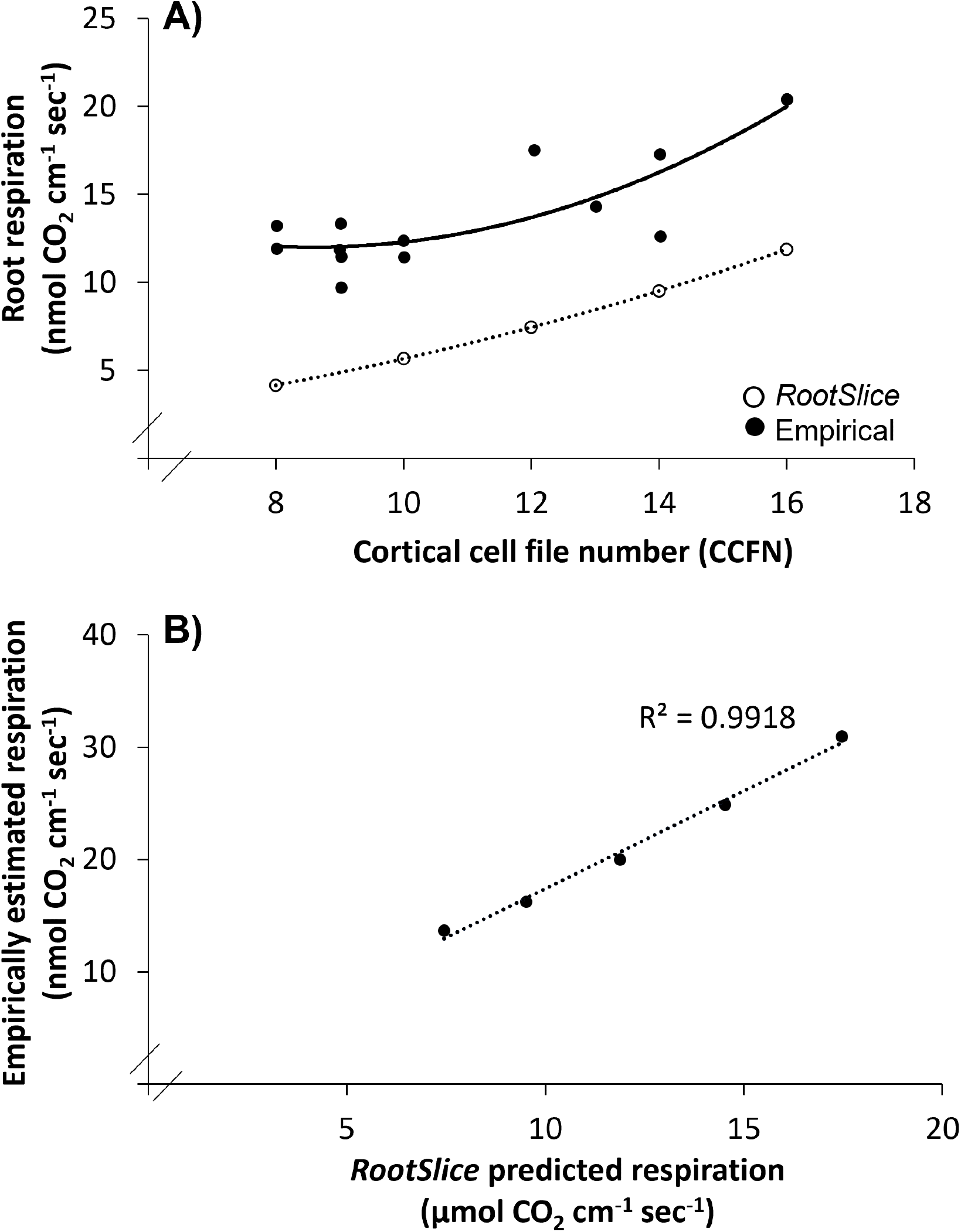
Comparing the *RootSlice* predicted effect of cortical cell file number on root respiration with empirical data (Chimungu *et al.,* 2014). A) *RootSlice* and empirical data showing an increase in root respiration with an increase in cortical cell file number, B) *RootSlice* predicted root respiration with changes in cortical cell file number compared to the empirical data.

### Case 4: Rhizoeconomic synergism between cortical aerenchyma and cell file number

*RootSlice* is capable of simulating phene interactions by showing interactive effects of cortical cell files and aerenchyma formation on root metabolic cost. With 12 cortical cell files, increasing aerenchyma formation to 40% of cortex volume was predicted to cause a ∼38 % decrease in respiration rate, ∼ 40% drop in nitrogen content, and ∼ 38 % decline in phosphorus content per unit root length (Figure 6C-E, 7B). On the other hand, the cortex with no aerenchyma and 8 cell files had ∼ 71% lower respiration rate, 78% less nitrogen content, and ∼78% reduced phosphorus content per unit root length, compared to that with 12 cell files (Figure 8D, E, 9A). Integrated root anatomical phenotypes with 8 cortical cell files and cortex volume with 40% aerenchyma led to the lowest rhizoeconomic variables compared to the phenotype with 20 cell files and no aerenchyma, with ∼83 % reduction in respiration rate, ∼88% lower nitrogen content, and ∼88 % decline in phosphorus per unit root length (Figure 10A, B, C). Aerenchyma formation and cortical cell files thus have an inverse rhizoeconomic relationship.

**Figure 10.**
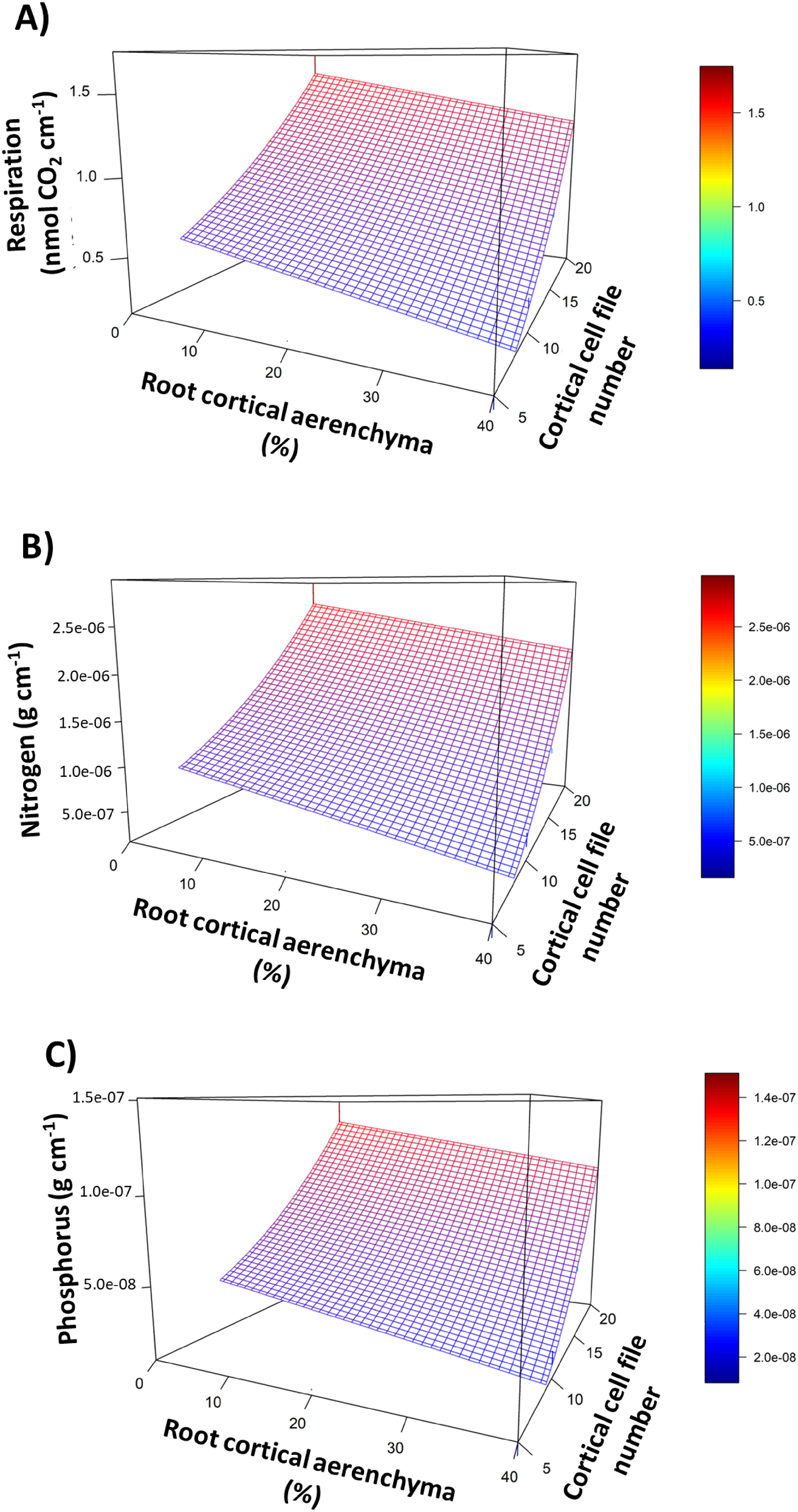
Response surface of the root respiration (A), root nitrogen content (B), and root phosphorus content (C) with root cortical aerenchyma and cortical cell file number. The quadratic function was found to be the best fit for the relationship of respiration, nitrogen, and phosphorus content with cortical aerenchyma and the number of cortical cell files.

### Case 5: Multiscale modeling integrating *RootSlice* with *OpenSimRoot* indicates that aerenchyma formation enhances the rhizoeconomic utility of fewer cortical cell files in improving shoot biomass performance under low nitrogen supply

Increased cortical cell files in nodal roots were predicted to reduce shoot biomass in varying nitrogen regimes (Figure 11A). The formation of cortical aerenchyma in the maturation zone up to a maximum of 20 and 40% of cortex volume in these phenotypes was predicted to improve shoot biomass performance (Figure 11 B, C). Root phenotypes with up to a maximum of 40% aerenchyma of cortex volume formation and 8 cortical cell files in the nodal roots outperformed all other phenotypes (Figure 11C). Reducing cortical cell files and increasing the formation of cortical aerenchyma in the nodal roots were predicted to increase the total length of the root system (Figure 11D, E, F). The formation of cortical aerenchyma decreases the total root carbon cost and increases the plant nitrogen content. The magnitude of this decline in root carbon cost and increase in nitrogen content varies with the number of cortical cell files and soil nitrogen supply (Supplementary Information 7). Phene synergism occurs when the benefits of a specific phene combination outweigh the additive effects, whereas antagonism causes the opposite. The additive responses (i.e., shoot biomass) for the twelve simulated anatomical phenotypes under low nitrogen supply were quantified based on the response of the reference phenotype (i.e., non-aerenchymatous nodal roots with 12 cortical cell files). Of these, the integrated phenotypes with 8 cortical cell files and up to a maximum of 20 or 40 % aerenchyma formation showed synergism with 0.28 - 0.73 g (i.e., 5.5 -12.7 %) greater shoot biomass than that expected by their additive response (Figure 12A, B, C). In contrast, the integrated phenotypes with 16 and 20 cortical cell files and forming up to the maximum of 20 or 40 % aerenchyma showed antagonism, with 0.2-0.63 g (i.e., 6.1 -17.4%) less shoot biomass than would be achieved by their expected additive effect (Figure 12B, C). Increasing the proportion of aerenchyma from the maximum of 20% to 40 % in the root increases its synergism with 8 cortical cell files and antagonism with 16 and 20 cortical cell files.

**Figure 11.**
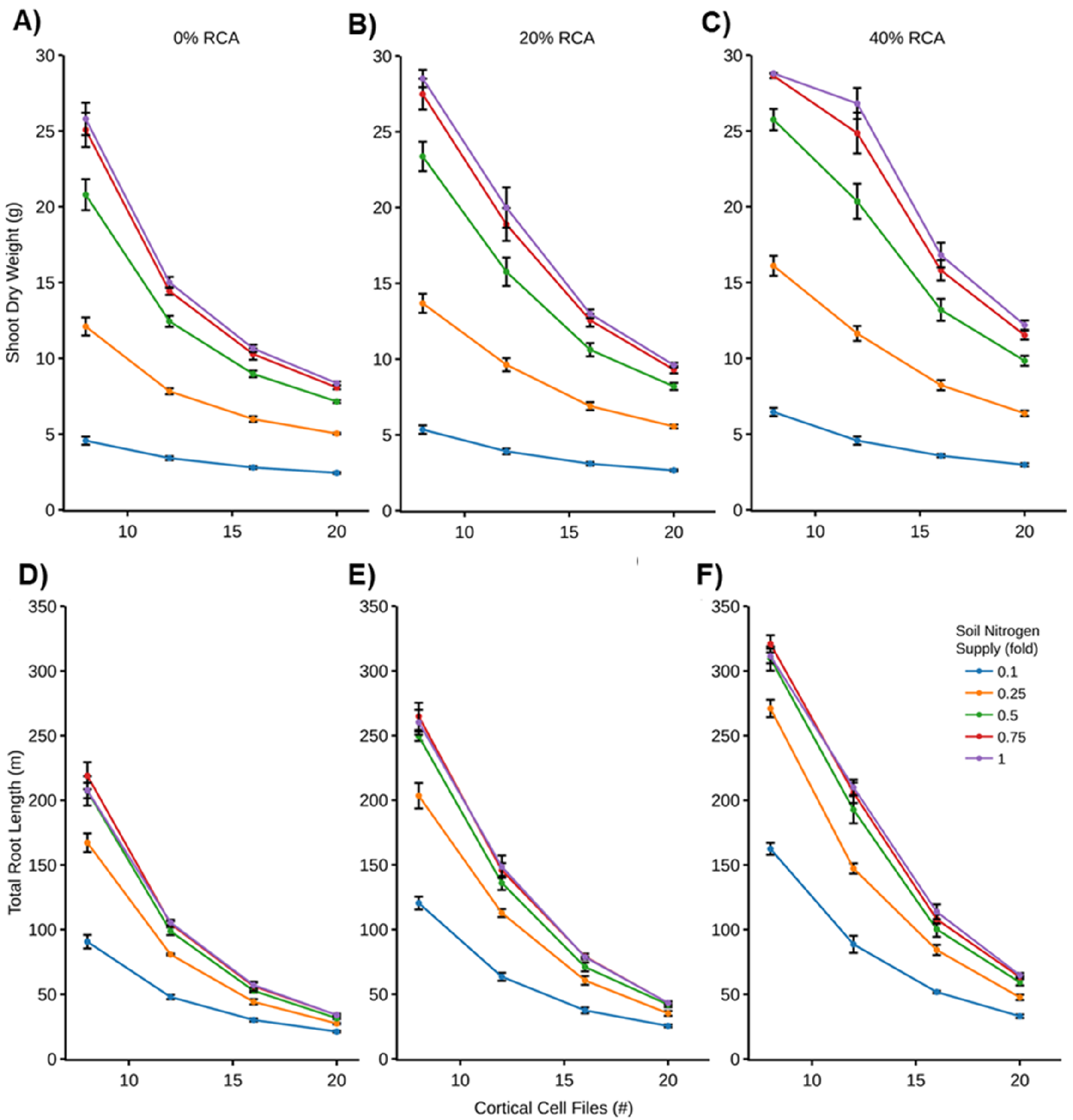
Shoot biomass (A-C), and total root length (D-F) of the maize root phenotypes over varied cortical cell files and aerenchyma formation under five levels of soil nitrogen supply.

**Figure 12.**
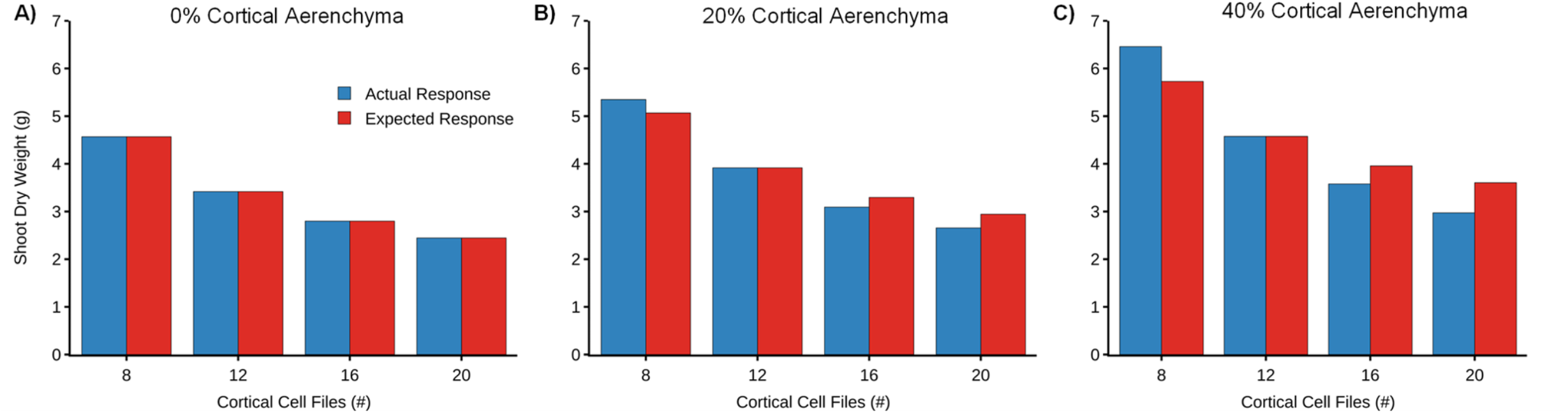
Quantifying interaction between root cortical aerenchyma and number of cortical cell files. Actual and expected (additive) responses (i.e., shoot biomass) of simulated phene combinations in maize under low nitrogen supply. Panel A, B, and C correspond to the different proportions of maximum cortical aerenchyma formed in the root (i.e., 0, 20, and 40%). Actual responses (blue) greater than, equal to, and less than the expected (red) respectively correspond to the synergistic, additive, and antagonistic relationship between the number of cortical cell files and the proportion of aerenchyma formation.

## Discussion

We present a new functional-structural model for root anatomy: *RootSlice*. We demonstrate the utility of this tool by evaluating the influence of diverse root anatomical phenotypes on underlying rhizoeconomics. To our knowledge, *RootSlice* is the first model capable of simulating three-dimensional multicellular root anatomies for diverse root segments. *RootSlice* captures different tissues and cell types with each cell consisting of distinct subcellular compartments, including cell wall, plasma membrane, cytosol, tonoplast, and vacuole. Currently, *RootSlice* is the only root anatomy model capable of simulating secondary thickening, which is a characteristic feature of mature dicotyledonous roots. Parameterization of *RootSlice* models involves subcellular and cellular morphometric data extracted from microscopic root images empirically acquired via e.g., manual sectioning, vibratome slices, laser ablation tomography, or μ-CT (Lynch *et al*., 2021; Strock *et al*., 2022). The model allows the specification of distinct resource demands for each subcellular compartment, including the differential concentration of nitrogen and phosphorus in vacuoles and cytosol. This enables the estimation of the total resource content per volume of the simulated root segment. Likewise, the mitochondrial density per unit volume of cytosol offers the estimation of the respiratory cost of the root segment with given anatomy. We demonstrate that the predicted rhizoeconomic variables for root segments corresponding to distinct developmental zones can be integrated as the temporal properties of growing roots in the functional-structural plant model *OpenSimRoot* to explore the utility of root anatomical phenotypes on plant performance in response to different edaphic and agroclimatic conditions. Such integration of *RootSlice* and *OpenSimRoot* models offers a genuine multiscale evaluation of root phenotypes.

With maize nodal roots as an example, we present five case studies involving vacuole expansion, the number of cortical cell files, and aerenchyma formation in this work. Focusing on the subcellular features of the model, in the first case study the model quantifies the role and metabolic consequences of vacuolar expansion in root elongation. In the second and third cases, the multicellular features of *RootSlice* confirmed the utility of fewer cortical cell files and a greater proportion of aerenchyma formation in reducing the metabolic cost of root segments. In the fourth case study, the combined utility of two anatomical phenes (i.e., number of cortical cell files and proportion of aerenchyma) in optimizing the rhizoeconomic cost of a root segment is evaluated. Extending the latter, the fifth case illustrates the multiscale rhizoeconomic evaluation of cortical cell file number and proportion of aerenchyma formation for plant performance under varying nitrogen availability.

Root cells can considerably expand in the elongation zone (Dhugga *et al*., 2004; Ivakov & Persson, 2013). The cell expansion is driven by turgor pressure, mostly provided by the expanding vacuole (Dünser *et al*., 2019; Scheuring *et al*., 2016). Each cell in the elongation zone expands until it reaches the mature zone where growth ceases and the secondary cell wall is deposited (Anderson *et al*., 2010; Marshall *et al*., 2012). *RootSlice* can simulate different root zones, starting from pre-elongation to maturation (Figure 2). *RootSlice* shows that majority of the cell expansion is because of expanding vacuoles while the volume of cytoplasm, plasma membrane, and the cell wall is increasing at a much slower rate and quantifies the consequences of vacuolar expansion on tissue metabolic costs (Figure 5).

*RootSlice* simulates cortical aerenchyma formation by following a programmed cell death algorithm (Buckner *et al*., 1998), which initiates the cell lysis in the mid cortex and then moves tangentially and radially. This pattern of aerenchyma formation has been seen in various species including wheat, sorghum, and maize (Drew *et al*., 2000; Promkhambut *et al*., 2011; Watkin *et al*., 1998). Some crop species, including rice, with extreme cortical aerenchyma formation occupying up to 80% of the root cortex, could also be captured using *RootSlice* (Figure 3G). To evaluate the functional implications of cortical aerenchyma, three different states of aerenchyma formation (i.e., 0, 20, and 40 %) in maize nodal roots were simulated. The model confirmed that aerenchyma formation led to cheaper roots by reducing the maintenance cost of the living cortical tissue (Figure 3H). Furthermore, simulations show that there is a ∼ 30 % reduction in protoplasmic volume, nitrogen content, phosphorus content, and mitochondrial density per cm^-3^ of cortex volume (Figure 6). In maize, roots with ∼ 20% aerenchyma formation were measured to have ∼25 to 57 % lower respiration than that without aerenchyma formation, thereby supporting the model prediction. Even though *RootSlice* prediction is slightly different in predicting the absolute decrease in nitrogen content and respiration due to aerenchyma formation, it accurately predicts the relative relationship of aerenchyma formation with respiration and nitrogen content (Figure 7).

The number of cortical cell files is known to influence various structural and functional aspects of the roots including its diameter, construction, and maintenance cost, radial resource flux, and penetration of hard soil (Chimungu *et al*., 2014; He *et al*., 2017; Vanhees *et al*., 2020). The impact of varying the number of cortical cell files on the rhizoeconomics of the maize nodal root segment was assessed using *RootSlice*. The model highlights that fewer cortical cell files reduce construction and maintenance costs. In maize, reduced cortical cell files were found to improve plant performance under drought stress (Chimungu *et al*., 2014) and low phosphorus availability (Galindo-Castañeda *et al*., 2018) compared to greater cortical cell files. This was attributed to their enhanced soil exploration as the result of lower root construction and maintenance of the roots.

Individual anatomical phenotypes have been shown to affect the metabolic cost of root tissue. However, the anatomical phenotype of a root segment is an emergent property of the states of multiple subtending phenes. Interaction among these phene states defines the functional and rhizoeconomic utility of the root anatomical phenotype and in turn, the soil exploration strategies of the root system (Klein *et al*., 2020). Reduced cortical cell file number and a greater proportion of aerenchyma individually decrease the root construction and maintenance cost. When combined, the rhizoeconomic cost of the integrated anatomical phenotype with 8 cortical cell files and 40 % aerenchyma led to even lower rhizoeconomic costs compared to their explicit individual costs. Understanding phene interactions are important as the potential utility of a phenotype cannot be predicted by its performance in an iso-phenic background (York *et al*., 2013). Gaining such insights by employing *in silico* approaches thus becomes vital for guiding crop improvement.

Coupling of multicellular tissue-scale *RootSlice*/*maize* with the functional structural plant model *OpenSimRoot/maize* is one of the novel features of this study, which allowed the multiscale evaluation of root phenotypes from a cell to the whole-plant scale. The integrated model confirmed previous reports that fewer cortical cell files and greater cortical aerenchyma individually improve the shoot biomass gain in maize over varying soil nitrogen supply (Figure 11) (Postma & Lynch, 2011; Saengwilai *et al*., 2014). Importantly, the model effectively underpinned the interactive roles of decreasing cortical cell files and increasing aerenchyma formation in reducing the overall root metabolic cost for better soil exploration and in turn, improving maize performance under suboptimal soil nitrogen availability. Roots with fewer cortical cell files have lower construction costs per unit length. Aerenchyma formation in such roots would thus enhance the rhizoeconomic utility of cortical cell files by lowering the proportion of living cost cells and their corresponding maintenance cost. The influence of aerenchyma formation on reducing root metabolic cost is mitigated by an increased number of cortical cell files. Unlike the number of cortical cell files, aerenchyma is a dynamic and evolving phene. The modeling highlights an increasingly synergistic relationship between the evolving aerenchyma formation (i.e., from 20 to 40%) and the cortex with 8 cell files for shoot biomass gain under low nitrogen availability. However, a similar increase in aerenchyma formation in the root phenotypes with 16 and 20 cortical cell files offers an increasingly antagonistic impact on shoot biomass in response to low nitrogen supply (Figure 12). This can be attributed to the parabolic variation in the cell size across the cortex. A greater number of cortical cell files skew the cell size distribution toward forming larger cells in the cortex (Supplementary Information 2). Thus, the cortex with a greater number of cell files would have a higher proportion of larger cells, which are relatively cheaper to maintain (Chimungu et al., 2014). This in turn reduces the root maintenance cost of per unit cortex volume. During the aerenchyma formation in the cortex with greater cell files, often the lysed cells are larger with lower metabolic cost. Therefore, the actual benefit of aerenchyma formation is lower than the expected benefit in the case of the increased number of cortical cell files due to the underlying proportional relationship between cell size and cortical cell files. On the other hand, the age-dependent increase in aerenchyma formation in the roots, highlights the spatiotemporally nature of these interactions between two or more spatial phenes. This temporal paradigm of phene interactions becomes important when applied in conjunction with adaptive plasticity of the root phenes, wherein a root phenotype would alter in response to edaphic stresses over time and in turn, its interaction with other phenes. Understanding such complex spatiotemporal interactions among root phenes certainly merits more attention and implementation of advanced *in silico*approaches.

The cortical cell files and size are key phenes contributing to root diameter. Assuming cortical cell size as a constant, varying the number of cortical cell files would alter the root diameter. A large root diameter is known to enhance the penetration ability of the root through hard soil layers (Materechera et al., 1992). Herein, the soil hardness is not simulated, which would have offered a trade-off for the altered rhizoeconomic cost thereby influencing the fitness landscape of the number of cortical cell files and root elongation. However, inherently thicker roots should be confused with ethylene-induced root thickening due to mechanical impedance which does not improve the penetration ability of roots through compacted soil layers (Vanhees *et al*., 2020). Likewise, the formation of root cortical aerenchyma comes with its own set of functional trade-offs, including reduced radial transport. Exploring the fitness landscape for the number of cortical cell files and the proportion of aerenchyma formation goes beyond the scope of this study, which is rather aimed at promoting the multiscale evaluation of root phenotypes to guide crop breeding.

Plants adapt to external stimuli through a cascade of regulatory mechanisms spanning different spatiotemporal scales. The *RootSlice* platform is a multicellular tissue scale model, which offers an initial platform to capture several dynamic phenomena. At a larger scale, the *RootSlice* model can be extended to capture processes such as root growth and cell expansion. Implementing root growth in the model would fundamentally require defining tissue anatomy as a time-dependent variable. This would allow the model to capture alterations in the root segment anatomy corresponding to cell division and differentiation over varying root zones with the formation of different morpho-anatomical features. With appropriate boundary conditions for the root segment, the model can be further extended to evaluate the influence of the evolving root anatomy on the biomechanical and kinematic properties of the root, respectively influencing its soil penetration ability and resource capture. Ultimately, such a root anatomical model can be envisaged to aid the root-centric ideotypes breeding for the reduced metabolic cost of soil exploration, improved soil resource capture, and better penetration of hard soil over an array of pedo-climatic conditions.

At the smaller scale, processes involving gene regulation, metabolic fluxes, hormonal signaling, resource uptake, transport, and storage can be implemented and assessed in the *RootSlice* model. Capturing biochemical dynamics would further add to the spatiotemporal complexity of the model. Coupling such a multicellular root growth model with models at larger or smaller spatiotemporal scales could require a significant amount of computational power. Given such challenges, the development of computational tools for integrating models at different spatiotemporal scales could become extremely helpful, enabling thorough in-silico evaluation of complex systems involving feedback mechanisms and phenotypic responses. The recent development of the polyglottic multi-scale model integration framework, named *Yggdrasil*, offers avenues for model integration across scales (Lang, 2019).

## Conclusion

Root anatomy plays an important role in soil resource acquisition but is complex. To facilitate the functional analysis of root anatomy and its variation, a new three-dimensional functional-structural model for root anatomy – *RootSlice*, was developed. The model accurately simulates root anatomical phenotypes in both cross and longitudinal sections. Subcellular compartmentation in the model allows the evaluation of subcellular phenes such as vacuolar size, which is shown to be important for cell elongation. In this study, the potential of *RootSlice* is shown by modeling the effect of cortical files, cortical aerenchyma, and their interaction on root metabolic cost. *RootSlice* coupling with *OpenSimRoot* shows that maize plants with the decreased cortical cell file number and increased root cortical aerenchyma would have the best performance under suboptimal nitrogen conditions as compared to all other phene combinations of these two phenes. Maize was used as an example in this study, however, *RootSlice* can be easily parameterized for other species as well, making it a flexible and generic model. Integration of molecular regulation models with a multicellular tissue-scale model like *RootSlice* opens new avenues for multiscale evaluation of root anatomical phenotypes on soil resource capture.

## Acknowledgments

IA, JPL and SA were supported by the Foundation for Food & Agriculture Research ‘Crops in Silico’ project (Grant ID 602757). The content of this publication is solely the responsibility of the authors and does not necessarily represent the official views of the Foundation for Food & Agriculture Research. JPL and JSS acknowledge support from the US Department of Energy ARPA-E Award DE-AR0000821. We acknowledge Jia Wu (Nanjing Agricultural University, China) for the initial development of the *RootSlice* model.

## Data Availability statement

Data will be available from the corresponding author on request.

## Supplemental Data

The following materials are available in the online version of this article.

**Supplementary Information 1.** Parameters used in different modules of *RootSlice*. Note some parameters were measured from the experiments conducted for this study.

**Supplementary Information 2.** Change in cell diameter among different cortical cell files. A) Representation of cortical cell files layout, B) quadratic curve fitted to model the change in cell diameter by cell file in different genotypes, each quadratic curve represents one genotype.

**Supplementary Information 3.** The algorithm of root cortical aerenchyma development in *RootSlice*. The close dotted line means the maximal lacuna area and the close curved line means the boundary of the zone. The green close curved line means middle zone, the yellow means down and up zone, and the blue means left and right zone. The arrows indicate the direction of Root Cortical Aerenchyma development in simulations.

**Supplementary Information 4.** Binary files for *RootSlice*, and sample input XML file.

**Supplementary Information 5.** *RootSlice* parameters simulating diverse root anatomies corresponding to different crop species and root classes are depicted in Figure 3.

**Supplementary Information 6.** *RootSlice* derived input parameters for *OpenSimRoot*.

**Supplementary Information 7.** Metabolic cost and plant nitrogen content of the root phenotypes differing in cortical cell file and aerenchyma formation predicted by the coupled *RootSlice-OpenSimRoot* model over varying soil nitrogen supply.

