## Supplementary Data for "*RootSlice* – a novel functional-structural model for root anatomical phenotypes"

**Supplementary Information 1.** Parameters used in different modules of *RootSlice. Note some parameters were measured from the experiments conducted for this study.*

| Parameters | | Value | Unit | Reference |
|---|---|---|---|---|
| **Geometry module** | | | | |
| | Cortical cell length | 75 - 350 | µm | Measured for this study |
| | Cortical cell diameter | 20 - 30 | µm | Schneider *et al.*, 2020 |
| | Cortical cell file number | 7 - 12 | | Schneider *et al.*, 2020 |
| | Cell wall thickness | 0.1 to 3.0 | µm | Schneider *et al.*, 2021; Lenochová *et al.*, 2009; measured for this study |
| | Tonoplast to plasma membrane distance | 0.2 - 3 | µm | Lenochová *et al.*, 2009; measured for this study |
| | Stele diameter | 200 - 1500 | µm | Schneider *et al.*, 2020 |
| **RCA module** | | | | |
| | RCA ratio | 0 - 80 | % | Schneider *et al.*, 2020; Mano *et al*. 2006 |
| | RCA number | 10 - 40 | | |
| **Rescource module** | | | | |
| | Cytoplasmic Protein N | 925 | mol $m^{-3}$ | Brown & Cartwright, 1953; Belton *et al*., 1985 |
| | Cytoplasmic Nucleic acid N | 57 | mol $m^{-3}$ | Brown & Cartwright, 1953; Belton *et al*., 1985; Close & Beadle, 2004 |
| | Cytoplasmic amino acid N | 57 | mol $m^{-3}$ | Brown & Cartwright, 1953; Belton *et al*., 1985; Close & Beadle, 2004 |
| | Cytoplasmic low weight organic compounds N | 57 | mol $m^{-3}$ | Brown & Cartwright, 1953; Belton *et al*., 1985; Close & Beadle, 2004 |
| | Cytoplasmic ammonium | 10 | mol $m^{-3}$ | Lee and Ratcliffe 1991 |
| | Cytoplasmic nitrate | 3.1 | mol $m^{-3}$ | Miller and Smith 1996 |
| | Vacuolar Protein N | 11 to 125 | mol $m^{-3}$ | Belton *et al.,* 1985; Mettler & Leonard, 1979; Wagner *et al*., 1981 |
| | Vacuolar Nucleic acid N | 92.5 | mol $m^{-3}$ | Belton *et al.,* 1985; Mettler & Leonard, 1979; Wagner *et al*., 1981; Close & Beadle, 2004 |
| | Vacuolar ammonium | 15 | mol $m^{-3}$ | Lee and Ratcliffe 1991 |
| | Vacuolar nitrate | 26 | mol $m^{-3}$ | Miller and Smith 1996 |
| | Nitrogen soluble fraction | 4 - 10 | % | Lee and Ratcliffe 1991 |
| | Cytoplasmic phosphate | 4.1 - 6.5 | mol $m^{-3}$ | Lee *et al*., 1990 |
| | Vacuolar phosphate | 1.0 - 9.0 | mol $m^{-3}$ | Lee *et al*., 1990 |
| | Mitochondrial density | 170* 1e+09 | $cm^{-3}$ cytoplasm | Close and Beadle 2004 |
| | Respiration per cytoplasm | 50 - 50000 | nmol $CO_2$ $cm^{-3}$ | measured for this study |
| | Cell wall density | 1.51 | g $cm^{-3}$ | Vatehová *et al*. 2016 |
| | Cell wall construction cost | 2.8 | MJ $g^{-1}$ cell wall | Estimated based on standard enthalpy |

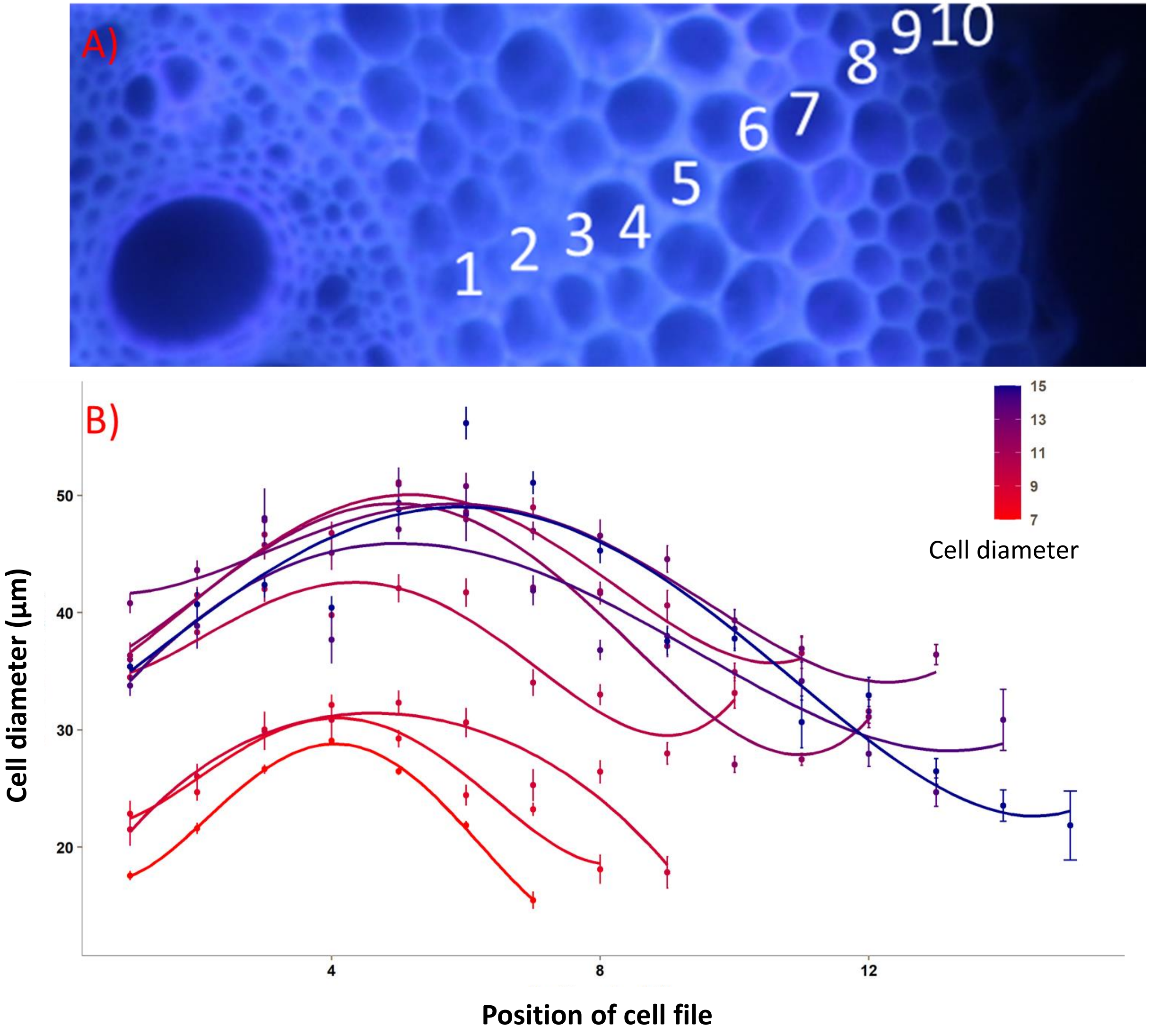


**Supplementary Information 2.** Change in cell diameter among different cell files. A) Representation of cortical cell files layout, B) quadratic curve fitted to model the change in cell diameter by cell file in different genotypes, each quadratic curve represents one genotype.

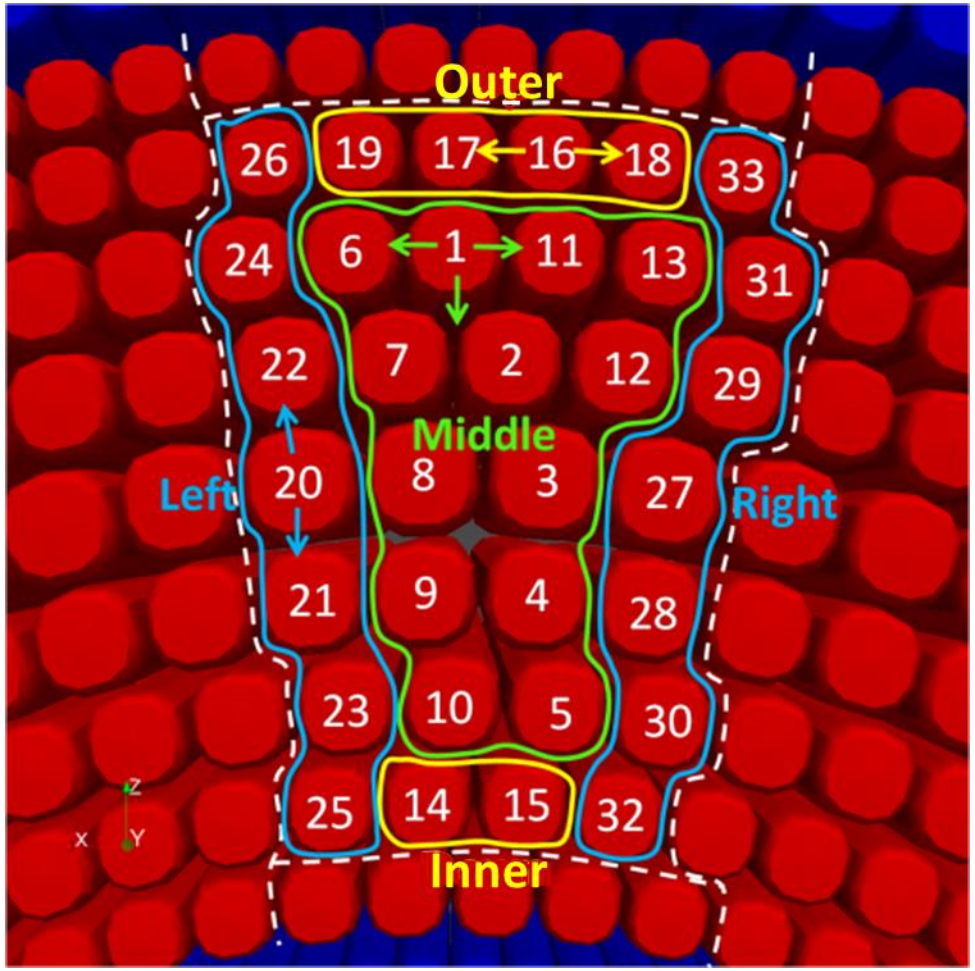


**Supplementary Information 3.** The algorithm of RCA development in *RootSlice*. The close dotted line means the maximal lacuna area and the close curved line means the boundary of the zone. The green close curved line means middle zone, the yellow means down and up zone, and the blue means left and right zone. The arrows indicate the direction of RCA development in simulations.

**Supplementary Information 4.** Binary files for the RootSlice and sample input XML file. The zip file has been uploaded to Zenodo.

Zenodo link: https://doi.org/10.5281/zenodo.6646845

**Supplementary Information 5. Simulation parameters used for simulating different crop species and root types in figure 3.**

| SPECIES (FIGURE A INDEX) | CWT (µm) | RCA | Cell length (µm) | CCFN | Stele diameter (µm) | CCD (µm) | Meta xylem number | Meta xylem radius (µm) |
|---|---|---|---|---|---|---|---|---|
| **Maize nodal root (3A)** | 2 | 0 | 100 | 8 | 400 | 30 | 10 | 25 |
| **Maize nodal root (3B)** | 2 | 0 | 100 | 12 | 400 | 30 | 10 | 25 |
| **Maize Braceroot (3C)** | 2 | 0 | 100 | 20 | 1000 | 15 | 45 | 25 |
| **Maize ancient nodal root (3D)** | 2 | 0 | 100 | 13 | 800 | 30 | 10 | 35 |
| **Pearl millet nodal root (3E)** | 2 | 0 | 100 | 8 | 280 | 32 | 4 | 25 |
| **Wheat nodal root (3F)** | 2 | 0 | 100 | 8 | 280 | 32 | 4 | 30 |
| **Wheat lateral root (3G)** | 2 | 0 | 100 | 4 | 160 | 60 | 4 | 5 |
| **Rice nodal root (3H)** | 2 | 65 | 100 | 12 | 280 | 16 | 4 | 30 |
| **Common bean basal root (3I)** | 2 | 0 | 100 | 6 | 120 | 50 | 17 | 30 |
| **Secondary growth in basal root (3J)** | 2 | 0 | 100 | 0 | 400 | 0 | 37 | 50 |

**Supplementary Information 6.** *RootSlice* derived input parameters for *OpenSimRoot*. The excel file has been uploaded to Zenodo.

Zenodo link: https://doi.org/10.5281/zenodo.6646845

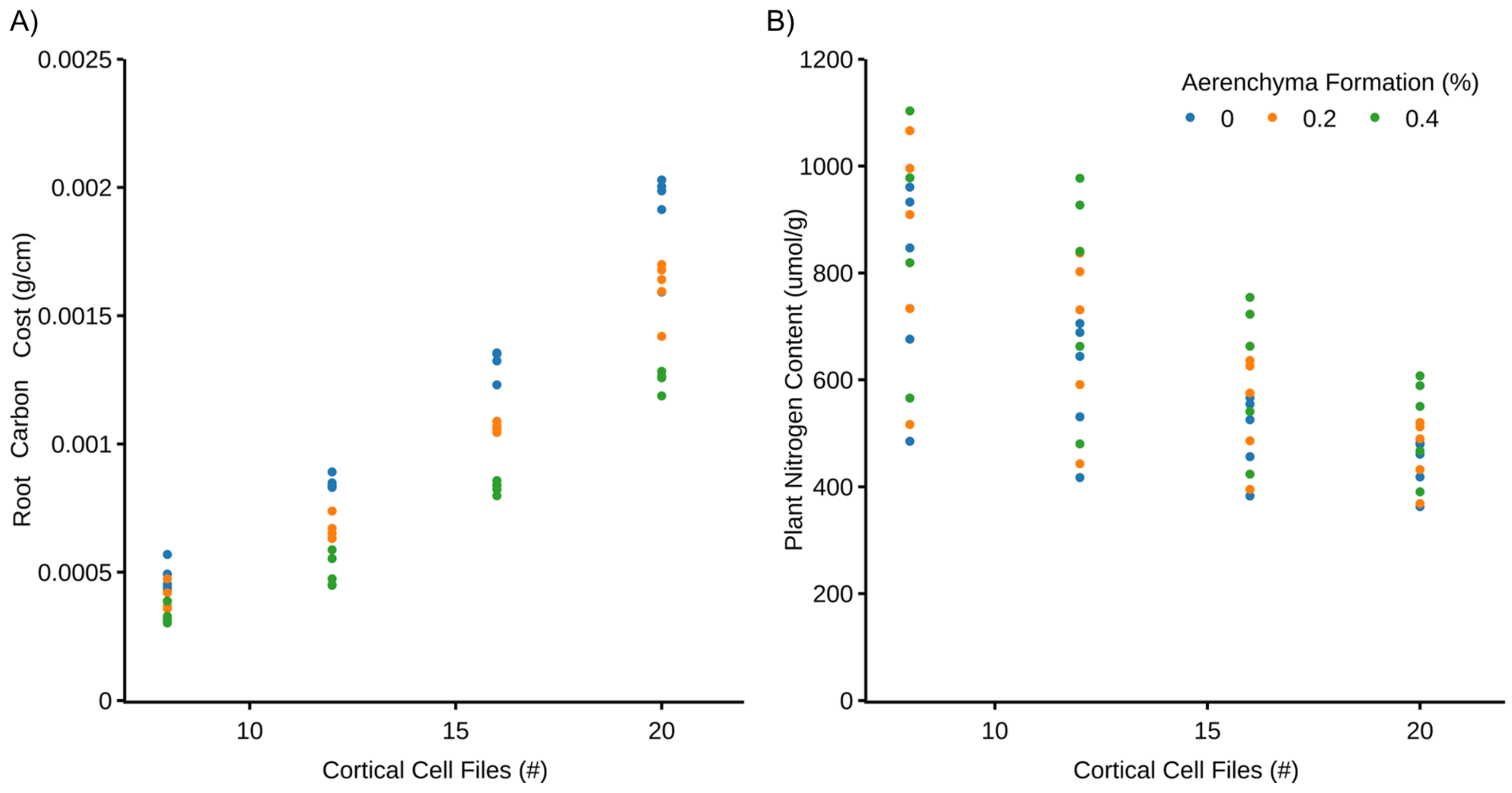


**Supplementary Information 7.** Root Carbon cost (A) and plant nitrogen content (B) of the root phenotypes differing in cortical cell file and aerenchyma formation predicted by the coupled *RootSlice-OpenSimRoot* model over varying soil nitrogen supply.
